# A barley MLA receptor is targeted by a non-ribosomal peptide effector of the necrotrophic spot blotch fungus for disease susceptibility

**DOI:** 10.1101/2023.12.13.571418

**Authors:** Yueqiang Leng, Florian Kümmel, Mingxia Zhao, István Molnár, Jaroslav Doležel, Elke Logemann, Petra Köchner, Pinggen Xi, Shengming Yang, Matthew J. Moscou, Jason D. Fiedler, Yang Du, Burkhard Steuernagel, Steven Meinhardt, Brian J. Steffenson, Paul Schulze-Lefert, Shaobin Zhong

**Affiliations:** Department of Plant Pathology, North Dakota State University, Fargo, ND 58108 USA; Department of Plant-Microbe Interactions, Max Planck Institute for Plant Breeding Research, Cologne 50829, Germany; Institute of Experimental Botany of the Czech Academy of Sciences, Centre of Plant Structural and Functional Genomics, Olomouc CZ-77900, Czech Republic; Cereal Crops Research Unit, Edward T. Schafer Agricultural Research Center, USDA-ARS, Fargo, ND 58102, USA; Department of Plant Pathology, University of Minnesota, St. Paul, MN 55108, USA; USDA-ARS Cereal Disease Laboratory, St. Paul, MN 55108, USA; Department of Computer Systems and Software Engineering, Valley City State University, Valley City, ND 58072, USA; John Innes Centre, Crop Genetics, Norwich Research Park, Norwich NR4 7UH UK; Cluster of Excellence on Plant Sciences, Max Planck Institute for Plant Breeding Research, Cologne 50829, Germany

## Abstract

The evolutionary history of plant interactions with necrotrophic pathogens that feed on dying host cells and their virulence mechanisms remains fragmentary. We have isolated the barley gene *Scs6*, which is required for the necrotrophic fungus *Bipolaris sorokiniana* isolate ND90Pr to cause spot blotch disease. *Scs6* is located at the disease resistance gene locus *Mildew locus a* (*Mla*) and encodes an intracellular nucleotide-binding leucine-rich repeat receptor (NLR). In transgenic barley, *Scs6* is sufficient to confer susceptibility to ND90Pr in accessions naturally lacking the receptor, resulting in infection-associated host cell death. Expression of *Scs6* in evolutionarily distant *Nicotiana benthamiana* reconstitutes a cell death response to an uncharacterized non-ribosomal peptide effector produced by ND90Pr-specific non-ribosomal peptide synthetases (NRPSs) encoded at the *VHv1* virulence locus. Our data suggest that the heat-resistant effector directly activates the SCS6 receptor. *Scs6* is an allelic variant of functionally diversified *Mla* resistance genes each conferring strain-specific immunity to barley powdery mildew isolates with a matching proteinaceous pathogen effector. Domain swaps between MLA and SCS6 NLRs and expression of the resulting hybrid proteins in *N. benthamiana* reveal that the SCS6 leucine-rich repeat domain is a specificity determinant for the NRPS-derived effector to activate the receptor. *Scs6* evolved after the divergence of barley from wheat and is maintained in several wild barley populations with an incidence of 8%, suggesting a beneficial function for the host. Evolution of the *bona fide* immune receptor SCS6 targeted by the NRPS-derived effector was key for the emergence of strain-specific spot blotch disease in domesticated barley.

## Introduction

Plants have evolved an innate immune system that is constantly challenged by a wide variety of microbial pathogens with different lifestyles, each of which has evolved different strategies to manipulate the host and establish virulence. Interactions between plants and biotrophic pathogens, which must retrieve nutrients from living host cells to proliferate, are often subject to co-evolution, with the pathogen restricted to a particular host species. The dynamics of these interactions are often driven by competing sets of co-evolving genes encoding plant immune receptors and pathogen effectors, the former being essential components for non-self-perception in the host and the latter being required for pathogen virulence (1). Despite recent advances, our understanding of the evolutionary history and dynamics of plant interactions with necrotrophic pathogens that kill and feed on dying host cells is less understood, even though these pathogens cause substantial economic damage in crops (2, 3).

Necrotrophic pathogens may have a wide or narrow host range. The molecular basis of host generalism is not well defined, but appears to be linked to the repertoire of secreted cell wall-degrading enzymes (3). Computational mining of pathogen genomes has revealed large arsenals of lineage- or species-specific effector proteins, often structurally related but with extreme divergence in their amino acid sequences (4–9). Experimental evidence shows that a subset of these effectors is required for virulence in necrotrophic pathogens with a narrow host range (2). Host-specialized necrotrophs often rely on proteinaceous or specialized metabolites that act as host-selective toxins (HSTs) to induce host cell death and promote infection. *Pyrenophora tritici-repentis* produces the proteinaceous ToxA effector, which targets the extracellular C-terminal domain of the wheat transmembrane protein TaNHL10, but susceptibility depends on wheat *Tsn1*, which encodes an intracellular hybrid protein consisting of an N-terminal S/T protein kinase fused to an NLR composed of nucleotide-binding (NB) and leucine-rich repeat (LRR) domains (10, 11). The necrotrophic *Parastagonospora nodorum* secretes the cysteine-rich proteinaceous effector SnTox1, which appears to directly target the plasma membrane-resident and wall-associated kinase (WAK) Snn1 for disease susceptibility in wheat (12, 13).

Chemically diverse metabolite effectors that act as HSTs have been identified in the fungal genera *Cochliobolus, Corynespora,* and *Periconia.* The susceptibility of sorghum to *Periconia circinata* depends on the *Pc* locus, which encodes a cluster of three tandemly repeated *NLR* genes and production of chlorinated peptide toxins by the pathogen, called peritoxins (14, 15). Loss of the central *NLR* results in loss of susceptibility to *P. circinata*, but it is unknown whether the toxin targets the NLR receptor directly or indirectly. The HC toxin of the causal agent of northern corn leaf spot, *Cochliobolus carbonum*, is a cyclic tetrapeptide and targets histone deacetylases of susceptible corn plants to establish infection (16, 17). *Cochliobolus victoriae* deploys victorin toxin, a mixture of ribosomally encoded but highly modified hexapeptides, to induce cell death and establish infection on *Vb*-containing oat genotypes (18). *Vb* is genetically inseparable from *Pc-2*, which mediates disease resistance to the biotrophic pathogen *Puccinia coronata*, but it remains unclear whether they are the same or two closely linked genes (19, 20). Victorin toxin is sufficient to induce cell death in several ‘non-host’ species, including about 1% of accessions of *Arabidopsis thaliana* (20–25). The NLR LOV1 in *A. thaliana* accession Cl-0 determines sensitivity to victorin, but also requires the thioredoxin *At*TRXh5, which contributes to salicylic acid-dependent defense through its denitrosylation activity on host proteins, including NPR1, the activator of systemic acquired resistance (26–28). Victorin binds to *At*TRXh5 and inhibits its activity. Since *At*TRXh5 binds to LOV1 in the absence of victorin, it is proposed that the receptor senses the toxin indirectly through victorin-mediated perturbation of *At*TRXh5 activity (23).

Isolate-specific disease resistance to biotrophic or hemibiotrophic pathogens is often conferred by intracellular plant NLRs that directly or indirectly sense the presence of pathogen effectors. This results in receptor oligomerization and resistosome formation, inducing immune signaling and termination of pathogen proliferation. Canonical plant NLRs consist of three domains, a variable N-terminal signaling domain, a central nucleotide-binding oligomerization (NOD) domain, followed by a C-terminal leucine-rich repeat region (LRR) (29). Most plant NLRs carry either a Toll-interleukin-1 receptor-like (TIR) domain or a coiled-coil (CC) domain at the N-terminus and are referred to as TNLs and CNLs, respectively (29, 30). The recognition specificity of sensor TNLs or CNLs is usually determined by their polymorphic LRR, whereas signaling NLRs become engaged in immune signaling initiated by sensor NLRs. CNL resistosomes integrate into host cell membranes and act as calcium-permeable channels that mediate Ca^2+^ influx, triggering immune signaling leading to host cell death (31–34). Sensor TNLs produce nucleotide-based second messengers that converge on the conserved EDS1 family to activate signaling/helper NLRs that carry a RESISTANCE TO POWDERY MILDEW 8 (RPW8)-CC domain (CC_R_) (31–38). Similar to sensor CNL resistosomes, activated signaling NLRs of *A. thaliana* have calcium-permeable channel activity (39). Ca^2+^ influx and the accumulation of reactive oxygen species are key events in immune signaling and are tightly linked to a regulated death of host cells at sites of attempted pathogen ingress, the so-called hypersensitive response (HR) (40–42). While the HR likely contributes to the termination of growth of biotrophic pathogens, it may promote the virulence of necrotrophs that retrieve nutrients from dying cells (43).

The necrotroph *Bipolaris sorokiniana* (*Bs*) (teleomorph *Cochliobolus sativus*) is the causal agent of a wide range of diseases in cereals, including leaf spot blotch, common root rot, seedling blight and kernel blight (44). Although *Bs* can infect a wide range of grass species, strain-specific variation in virulence among a worldwide collection of isolates has been identified based on differential infection responses on a panel of barley accessions, distinguishing four *Bs* pathotypes (45–47). Major genes or QTLs for spot blotch resistance/susceptibility have been identified in various barley genotypes depending on the *Bs* pathotype (48–54), but the dominant/recessive nature of each gene or QTL has yet to be determined in most cases. Recently, two wall-associated kinase genes, *Sbs1* and *Sbs2*, were isolated at the *Rcs5* locus, which confer susceptibility to spot blotch induced by the *Bs* isolate ND85F (55). Barley cultivar Bowman initially displayed moderate resistance to spot blotch when it was released in North Dakota, USA, in 1985 (56). Only six years later, Bowman and cultivars derived from Bowman showed hyper-susceptibility to a newly emerged isolate of spot blotch, named *Bs*_ND90Pr_ (57). This isolate belongs to *Bs* pathotype 2 and its high virulence on Bowman depends on the unique *VHv1* locus, which harbors a cluster of genes including two non-ribosomal peptide synthetases (NRPSs) (53, 58). Deletion of one of the two *NRPS* genes, termed *NPS1*, is sufficient to abolish the high virulence of *Bs*_ND90Pr_ on cultivar Bowman (58). We recently identified *Scs6* as the dominant gene needed for susceptibility to spot blotch caused by *Bs*_ND90Pr_ in Bowman and physically anchored the locus to a 125 kb genomic region overlapping with the *Mla* locus in the barley cv. Morex reference genome (59). Interestingly, the complex *Mla* locus is known to confer isolate-specific disease resistance to several foliar biotrophic pathogens, including the barley powdery mildew *Blumeria graminis* f. sp. *hordei* (*Bgh*), the stripe rust pathogen *Puccinia striiformis* and the hemibiotrophic blast pathogen *Magnaporthe oryzae* (60–64). The *Mla* locus harbors three NLR families, *Rgh1*, *Rgh2* and *Rgh3*, all of which encode CNL receptors (65). For several MLA CNL immune receptors belonging to the RGH1 family, cognate pathogen effector proteins, termed avirulence effectors, have been isolated and at least some bind directly to the corresponding receptor (66–69). Barley MLA immune receptors identified to date all belong to one of two MLA subfamilies from the RGH1 superfamily (63).

Here, we used chemical mutagenesis of the susceptible cultivar Bowman to identify several *Bs*_ND90Pr_ resistant mutants. A customized Mutant Chromosome Sequencing (MutChromSeq) (70) approach was then used to identify independent mutations in the susceptibility factor *Scs6*, which we show to be a naturally occurring *Mla* allele present in 16% of domesticated barley germplasm. We generated *Scs6* transgenic barley in accessions lacking the receptor to show that *Scs6* is sufficient to confer *Bs*_ND90Pr_ susceptibility. We collected intercellular washing fluids (IWFs) from Bowman leaves inoculated with wild-type *Bs*_ND90Pr_ or the *nps1 Bs*_ND90Pr_ mutant and show that the former IWF is necessary and sufficient to reconstitute a cell death response in *Scs6*-containing barley and in *N. benthamiana* transiently expressing *Scs6.* Domain swaps between the SCS6 CNL and the MLA1 or MLA6 barley powdery mildew immune receptors and expression of the resulting hybrid proteins in *N. benthamiana* revealed that the SCS6 LRR domain determines sensitivity to the NPS1-derived effector. We performed *Bs*_ND90Pr_ inoculation experiments with a collection of wild barley lines to show that *Scs6* is maintained in multiple geographically separated wild barley populations. Phylogenetic analysis suggests that *Scs6* is a *Hordeum*-specific innovation. We infer that SCS6 is a *bona fide* immune receptor that is directly targeted for disease susceptibility by the NPS1-derived effector of *Bs*_ND90Pr_.

## Results

### SCS6 is a naturally occurring variant of MLA subfamily 2 CNL receptors

To molecularly isolate *Scs6*, we applied the MutChromSeq approach (70) (**Fig. 1A**). We first mutagenized seeds of the susceptible barley cultivar Bowman with ethyl methanesulfonate (EMS; (71) and screened M_2_ families derived from approximately 1,500 M_1_ plants by inoculation of the seedlings with *B. sorokiniana* isolate ND90Pr (*Bs*_ND90Pr_) (**Fig. 1A**; Methods). A total of seven resistant M_2_ families (EMS14, EMS494, EMS621, EMS623, EMS787, EMS1317 and EMS1473) were identified, each characterized by drastically reduced cell death lesion formation in *Bs*_ND90Pr_-inoculated leaves compared to wild-type Bowman (**Fig. 1B**). Next, we flow-sorted chromosome 1H from five of the resistant EMS mutants and wild-type Bowman (**Fig. S1**), performed multiple displacement amplification (MDA) and BGISEQ-500 DNA sequencing of the 1H chromosomes (**Table S1**). We mapped sequence reads of each mutant line to the Bowman 1H assembly using the MutChromSeq pipeline and identified only one Bowman scaffold (scaffold_4918245 with a length = 23,130 bp) that was mutated in four mutant lines (EMS14, EMS621, EMS1317 and EMS1473) or deleted (the whole 23,130 bp sequence was missing) in one mutant (EMS494) (**Table S2**). The four mutant lines (EMS14, EMS621, EMS1317 and EMS1473) each carry different non-synonymous single nucleotide substitutions in a single gene (**Fig. 1C**). These substitutions are consistent with EMS alkylating activity on guanine residues and result in either premature stop codons or deduced single amino acid substitutions in the 5’ coding region of a candidate *Scs6* gene (**Fig. 1C**). Targeted genomic DNA resequencing of this gene, amplified by PCR from all seven mutant lines, validated the MutChromSeq analysis and identified two additional EMS mutant lines, EMS787 and EMS623, each carrying unique non-synonymous single nucleotide substitutions that resulted in a premature stop codon in the 5’ or a deduced single amino acid substitution in the 3’ coding region, respectively, making it likely that the corresponding wild-type gene is *Scs6* (**Fig. 1C**). The deduced protein of candidate *Scs6* consists of 959 amino acids with a tripartite domain organization typical of canonical CNL-type immune receptors, i.e., an N-terminal coiled-coil domain (CC), a central nucleotide-binding domain (NB), and C-terminal leucine-rich repeats (LRRs) (**Fig. 1C**). Protein sequence alignment with MLA/RGH1 variants found in multiple wild barley populations identified the candidate SCS6 as a novel member of the MLA receptor subfamily 2 (63). This subfamily differs from MLA subfamily 1 mainly by polymorphisms in the CC domain, but both subfamilies have an overall high protein sequence similarity of at least 88%.

**Figure 1.**
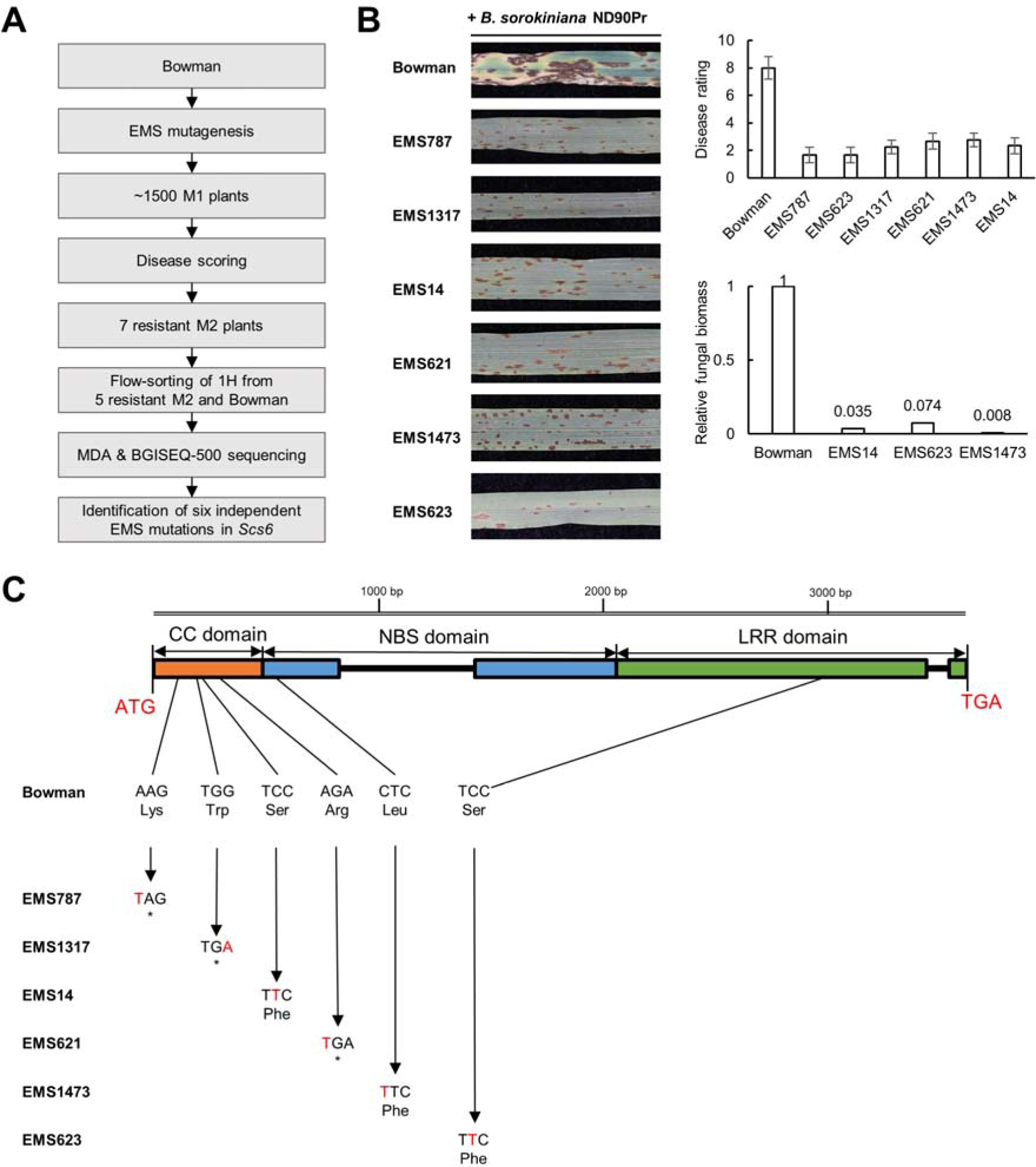
Identification of *Scs6* by MutChromSeq. (A) Workflow for MutChromSeq. (B) Infection responses, disease scorings, and quantification of fungal biomass in Bowman and six barley EMS M1 lines after inoculation with *Bipolaris sorokiniana* ND90Pr. Photos were taken at seven days after inoculation. The 1-9 rating scale of Fetch and Steffenson (96) was used to rate the spot blotch disease. Fungal biomass was quantified for Bowman and three EMS M1 lines using quantitative PCR. (C) Gene structure and EMS mutations in *Scs6*, a gene encoding a canonical coiled-coiled-type NLR (CNL).

### *Scs6* is necessary and sufficient to confer susceptibility to *Bs*_ND90Pr_ in barley

To further confirm that the candidate SCS6 confers susceptibility to *Bs*_ND90Pr_ in barley, we generated transgenic plant lines in *Bs*_ND90Pr_-resistant barley cultivar Golden Promise (GP) and barley line SxGP DH-47 (DH47) using two binary vectors that carry the coding sequence of candidate *Scs6* flanked either by the maize *Ubi* promotor and *NOS* terminator sequences or by 5’ and 3’ regulatory sequences of barley *Mla6*, respectively (**Fig. S2**). T_1_ progeny of T_0_ transgenic plants obtained from both GP and SxGP DH-47 genetic backgrounds showed a segregation pattern of strong susceptibility to *Bs*_ND90Pr_ that was dependent on the presence of *Scs6* transgene copies (**Fig. 2, Fig. S3 and Table S3**), validating that the candidate gene is *Scs6*. We conclude that *Scs6* is not only necessary for susceptibility to *Bs*_ND90Pr_ in cultivar Bowman but also sufficient to confer susceptibility to the fungal pathogen when introduced as transgene in both tested resistant barleys lacking the receptor.

**Figure 2.**
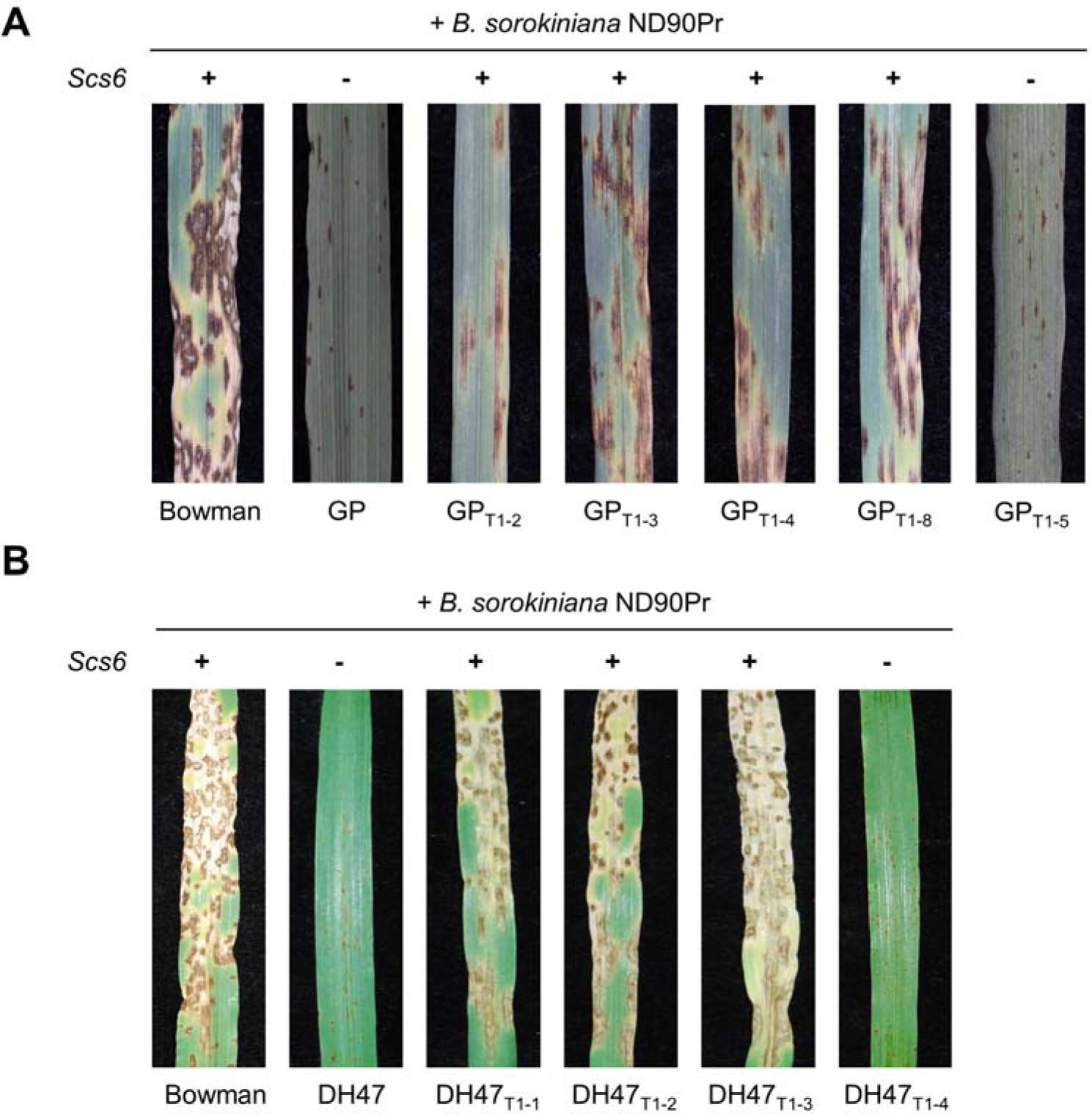
*Scs6* is necessary and sufficient to confer susceptibility to *Bipolaris sorokiniana* ND90Pr in barley. (A-B) Representative images of infection responses of Golden Promise (GP), SxGP DH47 (DH47) and derived transgenic *Scs6* T1 plants to *B. sorokiniana* ND90Pr, seven days after inoculation.

### Barley SCS6 is activated by a *Bs*_ND90Pr_ non-ribosomal peptide effector to induce cell death in barley and *N. benthamiana*

In previous studies, we identified two fungal genes in *Bs*_ND90Pr_ which encode a nonribosomal peptide synthetase (NRPS; *NPS1*) and a 4’-phosphopantetheinyl transferase (PPTase), respectively (57, 71). Both NPS1 and PPTase are necessary for *Bs*_ND90Pr_ to become virulent and induce necrotic lesions in Bowman leaves, and PPTase is required for activation of the NRPS enzyme (58, 72). We hypothesized that *Bs*_ND90Pr_ synthesizes and delivers a non-ribosomal peptide effector inside barley cells to induce SCS6-mediated cell death thereby facilitating its necrotrophic growth. We attempted to produce the effector by *in vitro* culture of *Bs*_ND90Pr_ in nutrient-limited media, but the fungal culture filtrates did not elicit necrotic symptoms after infiltration into Bowman leaves. We reasoned that the fungus might produce the effector during infection *in planta*. Therefore, we inoculated Bowman seedlings with wild-type *Bs*_ND90Pr_ and collected Intercellular Washing Fluid (IWF) from leaves seven days after the inoculation (denoted IWF_ND90Pr_, Methods). When IWF_ND90Pr_ was infiltrated into healthy leaves of Bowman, ND B112, and previously characterized double-haploid (DH) progeny derived from a cross between susceptible Bowman and resistant Culicuchima (59), only susceptible barley lines harboring *Scs6* developed necrotic lesions at the sites of IWF infiltration (**Fig. 3A, Fig. S4A and S4B**). This indicates that susceptibility to isolate *Bs*_ND90Pr_ and cell death activity of IWF_ND90Pr_ both depend on the presence of *Scs6*. IWF collected from barley leaves inoculated with the *Bs*_ND90Pr_ Δ*nps1* mutant (denoted IWF_Δ*nps1*_) failed to induce necrotic leaf lesions in *Scs6*-containing barley lines (**Fig. 3A**). Cell death activity of IWF_ND90Pr_ on Bowman was retained upon prolonged heat treatment of the IWF but lost after proteinase K incubation, consistent with an NRPS-derived effector (20 min 95 °C; **Fig. S4C**). Collectively, these results suggest that *Bs*_ND90Pr_ secretes a non-ribosomal peptide effector, that can be recovered by IWF extraction, to trigger *Scs6*-dependent cell death in barley.

**Figure 3.**
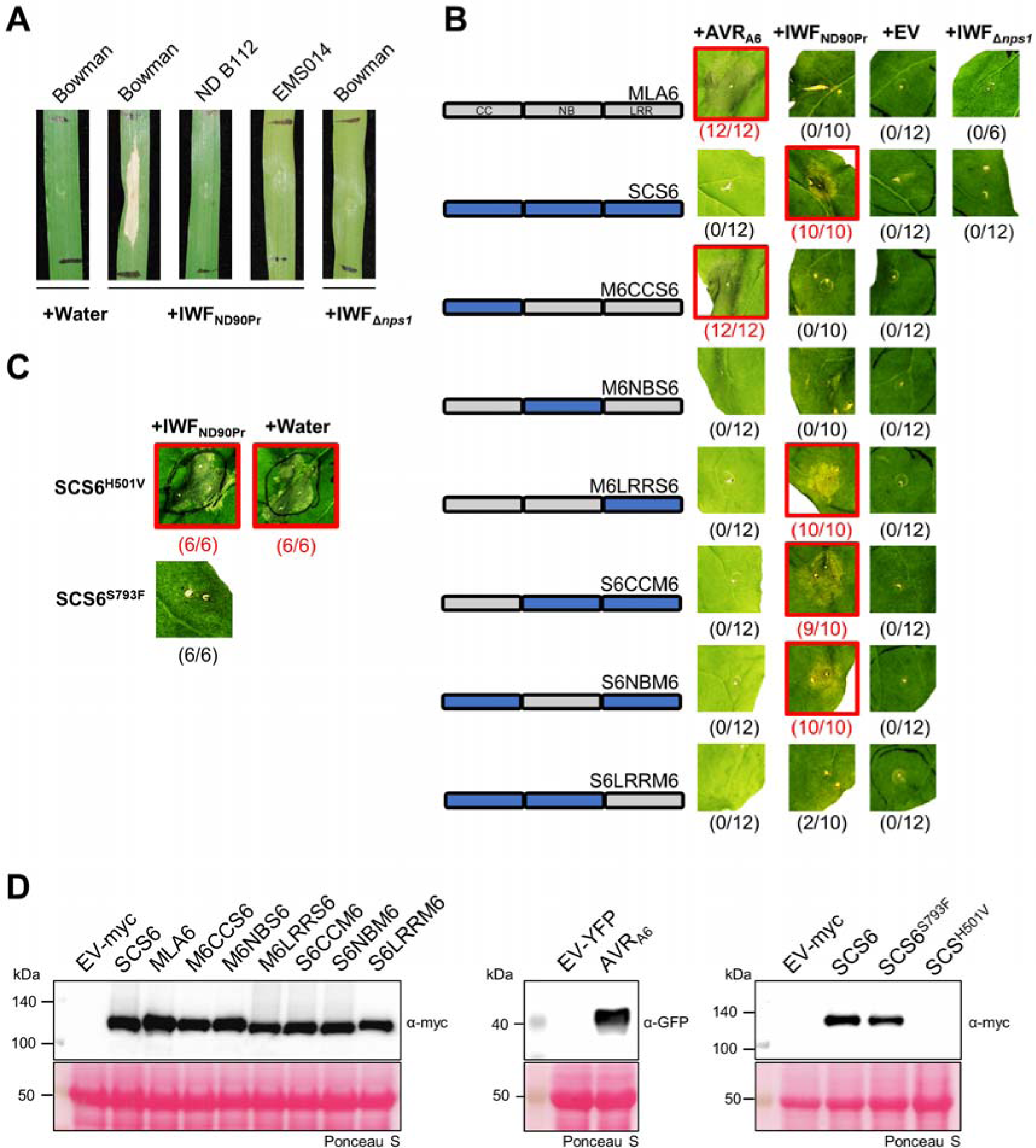
*Bipolaris sorokiniana* ND90Pr secretes an effector that activates Scs6 via its LRR region to cause cell death in barley and *Nicotiana benthamiana*. (A) Barley genotypes that express *Scs6* (Bowman) or negative control (ND B112) were infiltrated with intercellular washing fluid (IWF) that was isolated from Bowman leaves infected either with wild type *B. sorokiniana* ND90Pr (IWF_ND90Pr_) or mutant *B. sorokiniana* Δ*nps1* (IWF_Δ*nps1*_), as indicated. (B-C) *N. benthamiana* plants were transformed transiently, as indicated. Genes were fused in between the 35S promotor sequence and 4xmyc (receptors) or mYFP (AVR_A6_ without signal peptide) epitope sequences. Twenty-four hours after *Agrobacterium*-mediated gene delivery, IWF_ND90Pr_, IWF_Δ*nps1*_ or water was infiltrated, as indicated. Cell death phenotypes were assessed and documented at two or four days after agroinfiltration for IWF-triggered cell death or effector-triggered cell death, respectively. Representative pictures of at least six biological replicates (indicated in brackets) are shown and combinations that resulted in cell death are highlighted with a red box. (D) For determination of protein levels of receptor-4xMyc (approx. 114 kDa and AVR_A6_-mYFP (39 kDA) in *N. benthamiana*, leaf tissue was harvested two days post *Agrobacterium* infiltration.

To investigate whether SCS6 can serve as a target of the *Bs*_ND90Pr_-derived effector, we expressed the barley CNL in leaves of heterologous *Nicotiana benthamiana*, a dicotyledonous plant. We delivered wild-type *Scs6* or a *scs6* mutant *via Agrobacterium tumefaciens* infiltration. *Scs6* expression in *N. benthamiana* caused a rapid and robust induction of cell death after infiltration of IWF_ND90Pr_ but not IWF (**Fig. 3B**). Expression of *scs6* present in EMS mutant 623 (SCS6^S793F^) followed by IWF_ND90Pr_ infiltration also did not result in a cell death response. This is consistent with the finding that the EMS mutant 623 in barley is resistant to isolate *Bs*_ND90Pr_ (**Fig. 1C**), suggesting that the corresponding single amino acid substitution S793F in the SCS6 LRR domain renders the protein insensitive to the *Bs*_ND90Pr_-derived effector (**Fig. 3C**). Expression of a *Scs6* variant (SCS6^H501V^) resulting from a single amino acid substitution in the conserved MHD motif of the NB domain rendered SCS6 autoactive, i.e., SCS6^H501V^-mediated cell death in *N. benthamiana* occurred in the absence of IWF_ND90Pr_ (**Fig. 3C**). Equivalent substitutions in the MHD motif have been shown to result in autoactive MLA immune receptors triggering cell death *in planta* in the absence of matching *Bgh* avirulence effector proteins (73). Wild-type SCS6 and SCS6^S793F^ accumulated to similar steady-state levels in *N. benthamiana* leaf tissue (**Fig. 3D**). However, the auto-active SCS6^H501V^ variant was undetectable, presumably because little or no protein was produced due to immediate onset of cell death following *Agrobacterium*-mediated delivery of the corresponding gene construct (**Fig. 3D**). Taken together, these results demonstrate that barley *Scs6* expression in heterologous *N. benthamiana* is sufficient to recapitulate an IWF_ND90Pr_-dependent cell death.

### *Bs*_ND90Pr_-delivered effector specifically activates SCS6 *via* its LRR and NB domains

To further characterize SCS6-mediated cell death *in planta*, we constructed a series of hybrid receptors between SCS6 and MLA subfamily 1 immune receptors MLA6 or MLA1, guided by their shared modular domain architecture. The respective gene constructs were expressed in *N. benthamiana* following agroinfiltration and tested for their ability to induce cell death in the presence of matching *Bgh* avirulence effectors, AVR_A1_ or AVR_A6_, or IWF_ND90Pr_ or IWF_Δ*nps1*_ (**Fig. 3B**; **Fig. S5A**; (66). MLA1 and MLA6 were activated by cognate avirulence effectors AVR_A1_ and AVR_A6_, respectively, but not IWF_ND90Pr_, indicating that MLA recognition specificities for the proteinaceous and non-ribosomal peptide effectors are retained despite receptor overexpression. Hybrid receptors constructed through the exchange of the N-terminal CC domain of MLA1 or MLA6 with the corresponding sequence-diverged CC domain of SCS6 retained the ability to detect *Bgh* effectors AVR_A1_ or AVR_A6_, respectively (**Fig. 3B**; **Fig. S5A**). This is consistent with previous data showing that recognition specificities of MLA1 and MLA6 for the matching *Bgh* avirulence effectors are mainly determined by their polymorphic C-terminal LRRs (74). Similarly, SCS6 hybrids carrying the CC domain of either MLA1 or MLA6 retained the ability for cell death activation upon IWF_ND90Pr_ infiltration (**Fig. 3B**; **Fig. S5A**). This indicates that the CC domains of SCS6 and MLA1/MLA6 receptors are functionally interchangeable when mediating cell death in *N. benthamiana*, although the corresponding MLA subfamilies 1 and 2 are mainly differentiated by this polymorphic N-terminal CC module. Recognition of AVR_A1_ and AVR_A6_ by SCS6-MLA hybrids required the presence of both NB and LRR domains from MLA1/MLA6 receptors. A hybrid receptor carrying MLA6 CC and NB domains and the SCS6 LRR stimulated cell death upon IWF_ND90Pr_ infiltration, although cell death activity was slightly weaker compared to wild-type SCS6 (M6LRRS6; **Fig. 3B**). However, when the LRR of MLA1 was exchanged with the SCS6 LRR (M1LRRS6), the resulting hybrid was non-responsive to IWF_ND90Pr_ (**Fig. S5A**), indicating that both SCS6 NB and LRR domains are involved in SCS6 activation by the *Bs*_ND90Pr_ non-ribosomal peptide effector. All tested hybrid receptors accumulated to similar steady-state levels in *N. benthamiana* leaf tissue (**Fig. 3D**; **Fig. S5B**). These findings suggest that a *Bs*_ND90Pr_-released non-ribosomal peptide effector specifically activates SCS6 via its LRR and NB domains.

### *Scs6* susceptibility to spot blotch is common in barley

In nature, direct activation of SCS6-mediated cell death might be a strategy for the spot blotch pathogen to sustain its necrotrophic growth phase on susceptible barley. Therefore, we investigated the prevalence of *Scs6*-mediated susceptibility in domesticated and wild barley (**Fig. 4**, **Table S4 and S5**). We first performed *Bs*_ND90Pr_ inoculation experiments with 1,480 domesticated and 367 wild barley lines, the latter consisting of 318 accessions from the Wild Barley Diversity Collection (WBDC) and 49 additional *H. spontaneum* lines belonging to nine populations distributed throughout the Fertile Crescent (52, 75). We then conducted targeted DNA sequencing of *Mla* haplotypes to clarify whether susceptibility to *Bs*_ND90Pr_ spot blotch is strictly linked to the presence of *Scs6*, identified here as a member of MLA subfamily 2. The results showed that susceptibility to the fungal pathogen was invariably associated with the presence of *Scs6* identified in cv. Bowman (total of 269 accessions). Barley line FT153 was clearly susceptible to *Bs*_ND90Pr_ although previously only one MLA subfamily 1 variant was annotated at its *Mla* locus (*FT153-1*) (63), but the DNA sequencing of the corresponding genomic region detected a *Scs6* haplotype (*FT153-2*) that had escaped earlier analysis (63). Thirty-two wild barley accessions were susceptible to *Bs*_ND90Pr_ (**Figure 4A**). Based on targeted sequencing on twenty-one accessions and seven previously sequenced wild barley accessions (63), we confirmed that they all encode closely related SCS6 haplotypes (>97.90% protein sequence identity). FT170, for example, is highly susceptible and carries *FT170-1* as its sole subfamily 2 member, previously designated *Mla18-1* (63).

**Figure 4.**
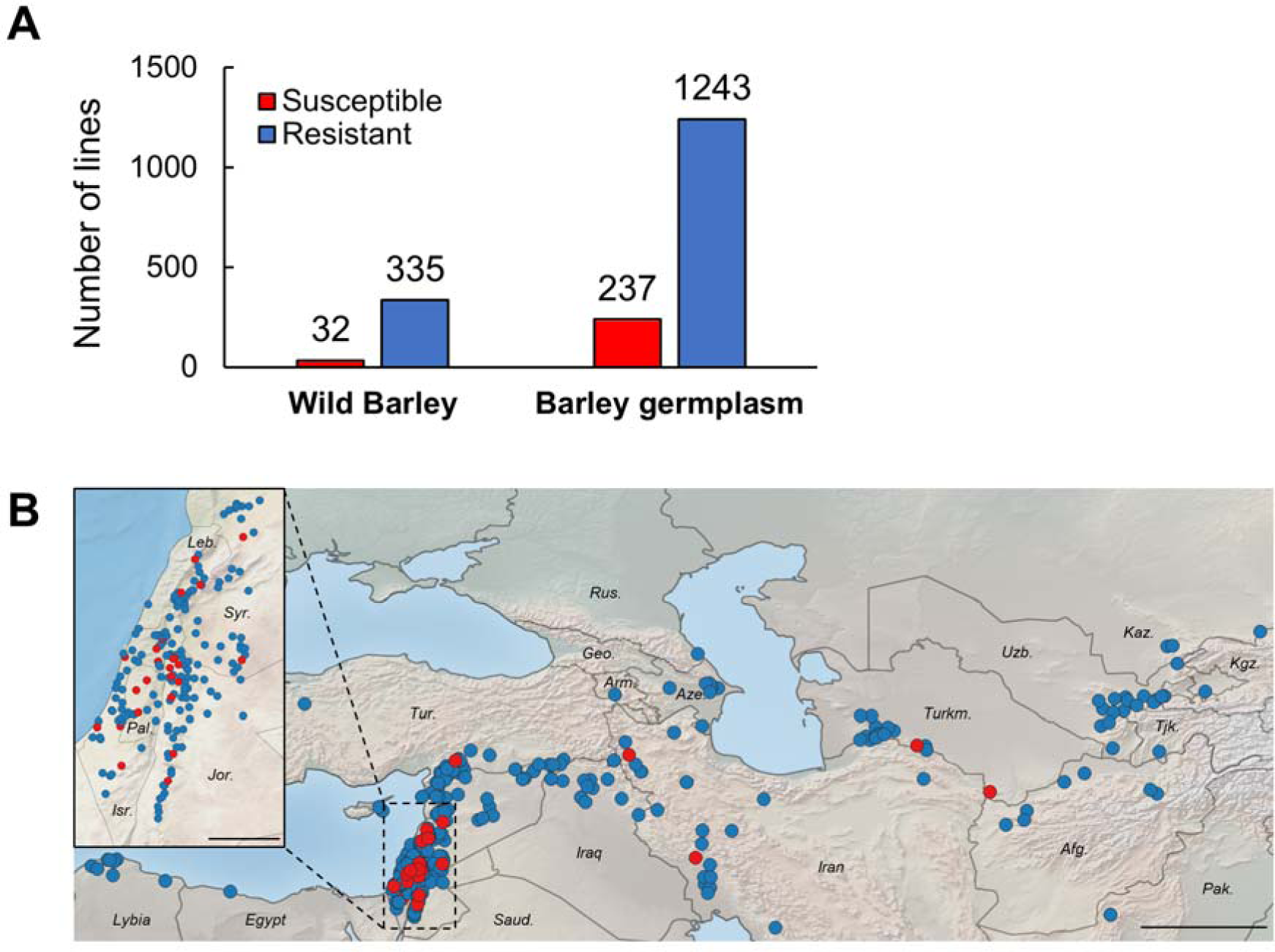
*Scs6* susceptibility to spot blotch is common in wild and cultivated barley. (A) Summary of inoculation experiments of wild barley (*Hordeum spontaneum*) accessions, including accessions from the Wild Barley Diversity Collection (WBDC; 52) and (63), and a panel of *Hordeum vulgare* germplasm with *Bipolaris sorokiniana* ND90Pr. (B) Geographic distribution of surveyed *Hordeum spontaneum* accessions. Susceptibility or resistance to *Bs*_ND90Pr_ is indicated in red or blue, respectively. Scale: 500 km (large map) and 100 km (map section on the left).

### The SCS6 receptor is likely a *Hordeum*-specific innovation

To investigate the evolutionary history of SCS6/MLA*-*mediated susceptibility to spot blotch, we curated a phylogenetic tree of all MLA variants found in wild and domesticated barley using neighbor-net analysis of full-length proteins. This revealed that SCS6 variants cluster within MLA subfamily 2 (**Fig. 5A**). In comparison to sequence divergence of individual MLA recognition specificities belonging to subfamily 1, sequence variation between SCS6 variants appear to be more limited although the corresponding accessions were sampled in distinct geographical regions and belong to different *H. vulgare subsp. spontaneum* populations (**Fig. 5A**). We examined an array of MLA subfamily 1 and subfamily 2 variants for sensitivity to IWF_ND90Pr_ in *N. benthamiana* and found that not only SCS6, but also subfamily 2 variants MLA16 and MLA18-1, can mediate effector-induced cell death and can therefore be considered SCS6 variants (**Fig. 5B**). However, sensitivity to the IWF was not shared among all MLA subfamily 2 members (e.g., MLA25, Sr50; **Fig. 5B and C**). This shows that there is natural genetic variation among all available MLA subfamily 2 members that accounts for their differential sensitivity to the *Bs*_ND90Pr_ NPS1-derived effector as well as susceptibility to the pathogen.

**Figure 5.**
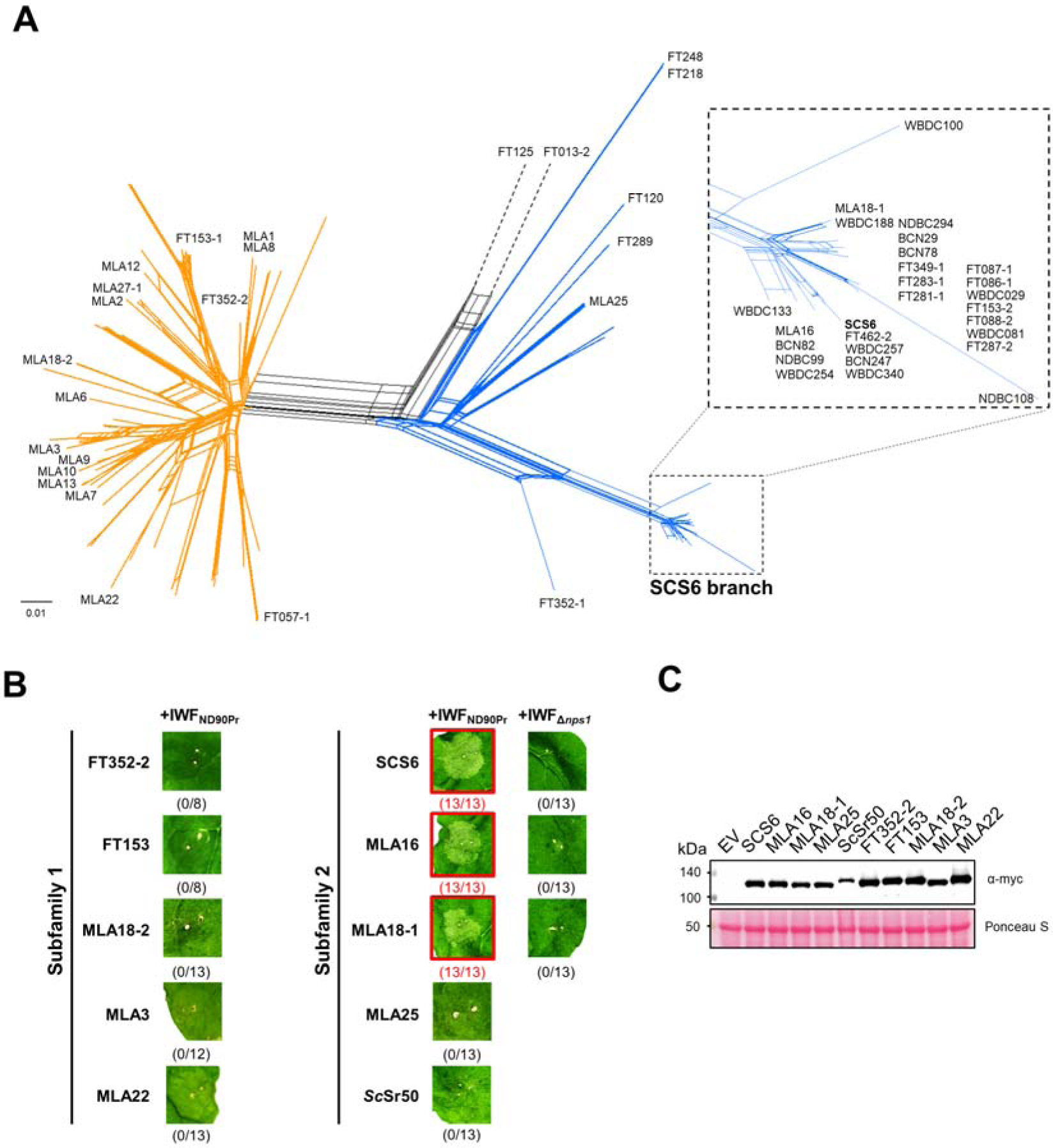
Diversity at the barley MLA locus underlies differential sensitivity to the *Bs*_ND90Pr_ NPS1-derived effector as well as susceptibility to spot blotch. (A) Neighbor-Net analysis of 114 MLA protein sequences including 28 previously identified MLA proteins from barley (64) 59 sequences from wild barley (63), as well as 27 sequences from wild or domesticated barley identified in this study. MLA subfamily 1 and MLA subfamily 2 members are represented using yellow or blue edges, respectively, based on (63) and Fig. S7. (B) *Nicotiana benthamiana* plants were transformed transiently, as indicated. Twenty-four hours after *Agrobacterium*-mediated gene delivery, IWF_ND90Pr_ or IWF_Δ*nps1*_ was infiltrated, as indicated. Representative pictures of at least eigth biological replicates (indicated in brackets) were taken two days after agroinfiltration and combinations that resulted in cell death are highlighted with a red box. OD_600_ of *A. tumefaciens* was set to 0.5, except for *Sc*Sr50, for which the OD_600_ was reduced to 0.2 to attenuate auto-activity. (C) Protein accumulation levels of expressed receptor-4xmyc constructs were determined by α-myc western blotting using total protein extracted from *N. benthamiana* leaves, one day post *agrobacterium-*infiltration.

We extended our aforementioned phylogenetic analysis, limited to *Hordeum* RGH1 variants, by including full-length proteins encoded by *Mla* orthologs or paralogs in other Triticeae species, including wheat (*Triticum*) and rye (*Secale*), and the wild grass *Dasypyrum villosum* (**Fig. S6**; (76)). MLA subfamilies 1 and 2 are mainly distinguished by their polymorphic CC domains (e.g., 65% identity and 81% similarity for MLA6 and SCS6 CC domains; (63)). The CC domains of some MLA haplotypes present in *D. villosum* can be assigned to MLA subfamily 1, while others are assigned to MLA subfamily 2 (**Fig. S7**), indicating that the differentiation of the CC domain occurred prior to the speciation of barley and *Dasypyrum villosum*, i.e., approximately 14.9 Mya, which predates the divergence of wheat and barley 8 Mya (77). Notably, we did not identify SCS6 homologs in other grass species, suggesting that SCS6 is likely a *Hordeum*-specific innovation. We performed statistical analysis on the coding sequences of *Scs6* variants, MLA subfamily 2 members from barley, and other *Mla* subfamily 2 members in the Triticeae to identify sites under positive selection. Strong signatures of positive selection in the LRR domain of Triticeae subfamily 2 members confirms and extends earlier analysis that MLA subfamily 2 includes resistance specificities against pathogens, e.g., Sr50 (**Fig. S8**; (63)). Limited positive selection detected among SCS6 variants may indicate that SCS6 evolves more slowly compared to the rapid evolution of MLA subfamily 1 members.

## Discussion

We have shown here that barley *Scs6* is necessary and sufficient to confer hyper-susceptibility to necrotrophic *Bs*_ND90Pr_. SCS6 is a canonical CNL encoded at the complex *Mla* locus on chromosome 1H, which harbors three highly dissimilar but physically linked *NLR* families, *Rgh1*, *Rgh2* and *Rgh3* (65, 78). All characterized disease resistance specificities at this locus have been assigned to the *Rgh1* family and a survey of wild barley revealed that *Rgh1* members are further sequence-diversified into two subfamilies, termed MLA subfamily 1 and subfamily 2 (63). Owing to the genomic head-to-head orientation of *Rgh2* and *Rgh3*, it has been proposed that they might act as paired NLRs against unknown pathogens (79). SCS6 shares 82% amino acid sequence identity with MLA6 and 28% and 24% sequence identity with RGH2 and RGH3, respectively, suggesting that a *bona fide* RGH1 member is needed for disease susceptibility of *Bs*_ND90Pr_. Expression of barley *Scs6*, but not barley *Mla1* or *Mla6*, in evolutionarily distant *N. benthamiana* reconstitutes a cell death response, specifically triggered by IWF collected from *Bs*_ND90Pr_ with an intact *VHv1* locus. Taken together with the capacity of autoactive SCS6^H501V^ to mediate cell death in the absence of a pathogen effector and the fact that all resistant EMS mutants carry mutations in *Scs6*, this indicates that SCS6 acts as a singleton NLR targeted by the NPS1-derived non-ribosomal peptide effector. The deduced function of SCS6 as a virulence target contrasts with characterized immune receptors encoded by *Rgh1*. In addition, only SCS6 is activated by a small molecule, whereas all other RGH1 members are activated upon sensing proteinaceous pathogen effectors to confer immunity (61, 66–69, 80, 81).

Drastically reduced fungal biomass on barley *scs6* leaves compared to wild-type *Scs6* Bowman following inoculation with wild-type *Bs*_ND90Pr_ suggests that *Scs6* is a virulence target for the fungus. As the reduced fungal biomass is tightly linked to loss of infection-associated host cell death on *scs6* mutants, it raises the possibility that *Scs6*-triggered signaling and/or cell death promotes the necrotrophic lifestyle of the spot blotch pathogen. Two deduced NRPSs are encoded at the *VHv1* locus in the *Bs*_ND90Pr_ genome and are unique to pathotype 2 strains (46, 58). Since deletion of one of the two *NRPS* genes at *VHv1* is sufficient to abolish high virulence of *Bs*_ND90Pr_ on cultivar Bowman (58), we conclude that a nonribosomally-encoded peptide effector produced by the fungus activates the SCS6 receptor.

Our data obtained with transgenic barley show that *Scs6* is the only host factor needed to render resistant barley cultivars lacking this CNL hyper-susceptible to *Bs*_ND90Pr_. This finding together with the observation that expression of barley *Scs6* is sufficient to reconstitute a cell death response in evolutionarily distant *N. benthamiana* in response to IWF_ND90Pr_ infiltration, strongly suggest that SCS6 is the direct virulence target for the NRPS-derived effector. Besides direct binding of pathogen avirulence effectors to the LRR domain, plant NLR receptors can also indirectly sense effector-mediated modifications in host proteins that serve as virulence targets (35, 82–84). In such an indirect activation model for SCS6 one would expect the formation of a pre-activation receptor complex through specific association with an unknown barley virulence target for the *Bs*_ND90Pr_-derived effector. As *Scs6* is shown here to be a lineage-specific innovation in barley (*Hordeum*), it seems unlikely that a pre-activation SCS6 complex can assemble in heterologous *N. benthamiana*, as this would imply an exceptional degree of evolutionary conservation of a hypothetical virulence target between dicotyledonous and monocotyledonous plants – species that diverged from each other approximately 140 Mya (85). Thus, our results contrast with the indirect recognition of the victorin toxin by the LOV1 CNL of *A. thaliana* through victorin-mediated disruption of *At*TRXh5 activity (23). In agreement with our conclusion, neither the expression of *LOV1* nor *AtTRXh5* alone in *N. benthamiana* leaves is sufficient to induce cell death after victorin infiltration (23). If *Vb*/*Pc-2* in oat is the same gene and encodes an NLR (19, 20), it will be interesting to test whether this receptor from the natural host of *C. victoriae* is directly or indirectly activated by victorin. Finally, the reconstitution of IWF-triggered and barley SCS6-dependent cell death in heterologous *N. benthamiana* suggests that the *Bs*_ND90Pr_ NPS1-derived effector can enter plant cells in the absence of pathogen infection structures and in the absence of a potential host species-specific surface receptor or transporter.

Similar to the proposed function of SCS6 as a direct virulence target for the *Bs*_ND90Pr_ NPS1-derived effector, experimental evidence strongly suggests that several other characterized barley RGH1 CNLs directly bind to proteinaceous avirulence effectors delivered by biotrophic *B. graminis* f sp *hordei* via the polymorphic LRR. These include MLA7, MLA10, MLA13, and MLA22, which respectively bind to sequence-diversified avirulence effectors AVR_A7_, AVR_A10_, AVR_A13_ and AVR_A22_ that share a common structural scaffold (66, 69, 81). Similar to SCS6, matching pairs of these MLA receptors and AVR_A_ effectors are necessary and sufficient to induce a cell death response in heterologous *N. benthamiana*. Additionally, the CNL receptor encoded by the stem rust resistance gene *Sr50* in wheat, an orthologue of barley *Rgh1* derived from rye chromosome 1R, assigned here to MLA subfamily 2, appears to bind directly to the stem rust effector AvrSr50 (67, 86). Collectively, this indicates that RGH1 CNLs have a propensity to interact directly with structurally distinct proteinaceous and even specialized non-ribosomal peptide effectors.

One of the EMS-induced mutants encodes a receptor variant with a single amino acid substitution in the LRR domain, SCS6^S793F^, which results in both loss of susceptibility to *Bs*_ND90Pr_ in barley and loss of cell death activity in response to IWF_ND90Pr_ infiltration in *N. benthamiana* (**Fig. 1 and Fig. 3**). Based on an AlphaFold2-generated SCS6 model, the residue S793 has an outward-facing side chain and is located on the concave side of the LRR. This, together with our observation that the SCS6 LRR domain is sufficient to confer IWF responsiveness to the corresponding MLA6 hybrid receptor M6LRRS6, corroborates an essential role of the SCS6 LRR as direct virulence target for the fungal-derived NRPS effector. In contrast, in the *Bs*_ND90Pr_-resistant barley mutants EMS14 and EMS1473, the deduced inward-facing receptor residues S73 and L183 are substituted by bulky phenylalanine, which is expected to destabilize the conformation of the CC and NB-ARC domains, respectively. If SCS6 functions similarly to sensor CNLs Sr35 and ZAR1 in wheat and Arabidopsis, then the latter two single amino acid substitutions in the SCS6 receptor might abolish the virulence activity of SCS6 by interfering with receptor oligomerization or Ca^2+^ pore formation after binding of the effector to the SCS6 LRR domain (31, 87). In addition to the LRR, the NB domain was found to contribute to the specific targeting of SCS6 by the peptide effector (**Fig. S5**), suggesting that the effector might interfere with NB and LRR interdomain interactions for receptor activation. A common activation mechanism for sensor CNLs and TNLs has recently been proposed on the basis of available cryo-EM structures of CNL and TNL resistosomes: Upon binding of proteinaceous avirulence effectors to the LRR domain, a steric clash with the central NB-ARC is triggered by the bound bulky protein effectors, inducing a conformational change of the NB-ARC domain followed by the exchange of ADP for ATP (88). However, NRPS-generated peptides are significantly smaller than known proteinaceous avirulence effectors, generally < 10 amino acids in length (89). Consequently, future work will address the question of how a much smaller NRPS-derived effector can both bind to the SCS6 LRR and induce a steric clash with the central NB domain.

Although *B. sorokiniana* isolates are typically generalists that can infect a wide range of Triticeae species, including wheat, the isolate *Bs*_ND90Pr_ is specialized to barley hosts. This is consistent with our finding that *Scs6* alleles were not detected in wheat or wheat progenitors, suggesting that *Scs6* might be a *Hordeum*-specific innovation that evolved after the divergence of the genera *Triticum* and *Hordeum* less than 8 Mya (90). This could explain why *Bs*_ND90Pr_ confers hyper-susceptibility only on *Scs6* barley genotypes, raising the possibility that *Bs* pathotype 2 acquired its unique *VHv1* virulence gene cluster during interactions with *Hordeum* hosts. However, whether *VHv1* of *Bs*_ND90Pr_ evolved as a post-domestication event in agricultural environments or in wild barley pathogen populations and subsequently spread to North America remains to be clarified.

All characterized *Mla* powdery mildew disease resistance specificities in barley belong to *Mla* subfamily 1, whereas no disease resistance function has yet been assigned to barley *Mla* subfamily 2, which includes *Scs6*. Extensive data support the notion that functional diversification of MLA subfamily 1 members is driven by a co-evolutionary arms race with the genetically highly diverse biotrophic *Bgh* pathogen (63, 91–93). Compared to MLA subfamily 1 members, our analysis of positive selection among subfamily 2 members indicates much less functional diversification, which is particularly striking among naturally occurring *Scs6* variants. We have shown here that SCS6 is maintained in several wild barley populations with an incidence of about 8%, strongly suggesting a beneficial function for the host. This widespread occurrence and the ability of autoactive SCS6^H501V^ to trigger cell death in the absence of a pathogen effector, makes it likely that the SCS6 CNL confers immunity against an unknown biotrophic or hemi-biotrophic pathogen endemic to barley populations in the Fertile Crescent. The postulated pathogen may not engage in a rapid co-evolutionary arms race with extant *Hordeum spontaneum* germplasm.

The hyper-virulent *Bs*_ND90Pr_ isolate emerged five years after barley cultivar Bowman was introduced in North Dakota in 1985. Unexpectedly, our pathotyping survey shows that *Scs6*-dependent susceptibility to *Bs_ND90Pr_* is twice as high in domesticated barley as in wild barley populations (16% and 8%, respectively). Domestication and breeding for disease resistance in barley may have inadvertently resulted in the co-enrichment of *Scs6*-dependent disease susceptibility to *Bs_ND90Pr_*, probably due to linkage drag from another disease resistance gene on barley chromosome 1H. Recently, the *Pyrenophora teres* f. *maculata* susceptibility factor *Spm1* was mapped to the *Mla* locus in the cultivar Baudin (94). Although it remains to be tested whether *Spm1* is also a member of the *Rgh1* family, our results demonstrate that the evolution of allelic variants of a single *R* gene is shaped by contrasting selective pressures exerted by multiple pathogens with different lifestyles. Elucidating the molecular principles underlying SCS6 activation by the NPS1-derived effector is likely to be of broader importance, as this could aid future development and deployment of synthetic NLR receptors in crops that are less vulnerable to manipulation by economically important necrotrophic pathogens.

## Materials and Methods

### Plant materials and generation of EMS mutant population

The barley cv. Bowman carrying *Scs6* (1) was used to generate mutant lines that were resistant to spot blotch caused by *B. sorokiniana* isolate ND90Pr. The mutagenesis procedure was performed according to (2) with some modifications. Approximately 2,000 seeds of barley cv. Bowman were presoaked in 300 ml of phosphate buffer (0.05 M, pH 8.0) for 8 hours at room temperature with gentle agitation. Then, the seeds were treated in 0.3% (v/v) ethylmethane sulfonate (EMS) in phosphate buffer for 16 hours at room temperature. Treated seeds were rinsed with water for 1 min and sown in pots immediately. The M_1_ plants were grown in the greenhouse at 20 to 24 °C under supplemental fluorescent lighting with a 16/8-h day/night cycle. Spikes were harvested separately from individual M_1_ plants. Approximately 20 M_2_ seedlings from each M_1_ plant were screened for spot blotch resistance using isolate ND90Pr following the procedures described by (3). Bowman was included as a positive control for susceptibility and ND5883 and NDB112 as positive controls for resistance. Resistant M_2_ seedlings were selected and propagated by selfing to develop homozygous M_3_ mutant lines, which were further confirmed for resistance to ND90Pr and then used for MutChromSeq analysis. Cultivated barley accessions from the USDA National Small Grains Collection (4) and the Wild Barley Diversity Collection (WBDC) accessions (5) were also screened against isolate ND90Pr and used in the *Scs6* gene diversity study.

### Fungal isolate, spot blotch phenotyping, Intercellular Washing Fluid (IWF) extraction and relative fungal biomass quantification

The pathotype 2 isolate ND90Pr of *B. sorokiniana* was used for phenotyping throughout this research. V8 PDA (150 mL of V8 juice, 850 of mL H_2_O, 10 g of PDA, 10 g of agar and 3 g of CaCO_3_) was used to culture *Bs*_ND90Pr_ under the conditions of 14 h of light and 10 h of darkness. Spore suspension containing 2×10^3^ conidia/ml was prepared and sprayed on seedlings with the second leaves fully expanded (12–14 days after planting). Inoculated plants were incubated in a humidity chamber for 18–24 hours and then transferred into the same greenhouse room. Disease ratings were conducted at 7 days post-inoculation using the 1–9 rating scale of (3).

To prepare the IWF, barley cv. Bowman was inoculated with *Bs_ND90Pr_*or *Bs*_ND90Pr_ Δ*nps1* as described above, and leaves were harvested 7 days after inoculation. Harvested leaves were cut into fragments of about 1 inch in length, and leaf fragments were submerged into distilled water in the beaker. The beaker was then set into a vacuum chamber and vacuumed for 30 minutes. Then, leaf fragments were surface-dried and transferred into 50-ml centrifuge tubes, which were centrifuged at 3900 rpm for 30 minutes. Finally, IWFs were harvested from the bottom of each centrifuge tubes, confirmed on barley cv. Bowman seedlings by infiltration, and stored at −20 °C for further use.

To quantify the fungal biomass, DNA was extracted from the leaves harvested at 7 days after pathogen inoculation using a DNeasy Plant Mini Kit (Qiagen, Germany). Subsequently, 50 ng of each DNA sample were used for quantitative real time PCR (qPCR), which was performed using the ITS region as fungal target and the actin gene of barley as reference. Real-time PCR was performed as described by (6). The ITS Ct values were normalized using the barley actin gene, and the relative gene copy number of ITS was calculated according to the 2^-ΔΔCT^ method (7). The relative quantity of fungal biomass was calculated using barley cv. Bowman leaves inoculated with wild type isolate ND90Pr as a control.

### Flow sorting of barley chromosomes and preparation of DNA for sequencing

Suspensions of mitotic metaphase chromosomes were prepared from root tips of barley cv. Bowman carrying SCS6 and its five EMS mutants following (8). Briefly, root-tip cells were synchronized using hydroxyurea, accumulated in metaphase using amiprohos-methyl and fixed by formaldehyde. Intact chromosomes were released by mechanical homogenization of 100 root tips in 600 µL ice-cold LB01 buffer (9). GAA microsatellites on the isolated chromosomes were labelled by fluorescence *in situ* hybridization in suspension (FISHIS) using 5’-FITC-GAA7-FITC-3’ oligonucleotides (Sigma, Saint Louis, USA) according to (10) and chromosomal DNA was stained by DAPI (4’,6-diamidino 2-phenylindole) at 2 µg/mL. Bivariate chromosome analysis and sorting was done using a FACSAria II SORP flow cytometer and sorter (Becton Dickinson Immunocytometry Systems, San José, USA). Sort window delimiting the population of chromosome 1H was setup on a dot-plot FITC-A vs. DAPI-A and 55,000–70,000 copies of 1H chromosomes were sorted from each sample at rates of 1,500–2,000 particles per second into PCR tubes containing 40 μL sterile deionized water. Chromosome content of flow-sorted fractions was checked by microscopic observation of 1,500–2,000 chromosomes flow sorted into 10 μL drop of PRINS buffer containing 2.5% sucrose (11) on a microscopic slide. Air-dried chromosomes were labelled by FISH with a probe for GAA microsatellite according to (12). In order to determine chromosome content and the purity, which was expressed as percent of 1H in the sorted fraction, at least 100 chromosomes in each sorted sample were classified following the molecular karyotype of barley (12). The samples of flow-sorted chromosomes 1H were treated with proteinase K, after which their DNA was amplified by multiple displacement amplification (MDA) (**Table S1**) using an Illustra GenomiPhi V2 DNA Amplification Kit (GE Healthcare, Chalfont St. Giles, United Kingdom) as described by (13). The DNA samples were sequenced by BGI using BGISEQ-500 (Cambridge, MA) to generate 100-bp paired-end (PE) reads.

### MutChromSeq

Raw sequencing data from flow-sorted chromosome 1H of the wild type and EMS mutants were quality-trimmed using Trimmomatic (14). The Bowman 1H chromosome sequencing data was assembled using ABySS 2.0 (15, 16) and was masked for repeats using RepeatMasker (http://repeatmasker.org). Sequence reads from EMS mutants were aligned to the repeats-masked Bowman 1H assembly using software BWA (17). The reads-aligned bam files were further processed using SAMtools 0.1.19 (17) following parameters suggested by (18). The resulting pileup formatted files for wild type and EMS mutants were used as the inputs analyzed by Pileup2XML.jar (https://github.com/steuernb/MutChromSeq). Finally, MutChromSeq.jar (https://github.com/steuernb/MutChromSeq) was executed to identify the candidate contigs with mutations in EMS mutants analyzed. All mutations were manually validated using Integrative Genomics Viewer software (IGV, version 2.5.2, (19)).

### Identification of the candidate gene for *Scs6*

Gene annotation for the MutChromSeq-identified contig with mutations in EMS mutants was performed by FGENESH (20). The genomic structure of *Scs6* was confirmed by PCR sequencing using both genomic DNA and cDNA as templates and primers listed in **Table S6**. *Scs6* was amplified by PCR from the five EMS mutants used in MutChromSeq and three additional EMS mutants with primer pair SCS6-F2/SCS6-R17 (**Table S6**).

### Binary vector construction and *Agrobacterium-*mediated transformation of barley

To determine the function of *Scs6*, two expression vectors of *Scs6* were constructed and used to transform Golden Promise and SxGP DH-47 (DH47), which are resistant to isolate ND90Pr, using the *Agrobacterium*-mediated transformation method. The whole coding sequence (CDS) of *Scs6* was synthesized by GenScript (Piscataway, NJ) and inserted between the *Spe*I and *Bsr*GI restriction sites of the binary vector pANIC12A (21), producing a new construct pANIC12A-*Scs6* with the *Scs6* gene driven by a *Ubi* promoter and stopped by a *NOS* terminator. Another binary vector (22) (pBract202-pMla6-*Scs6*-tMla6, **Figure S2**) was constructed, which carries the coding sequence of candidate *Scs6* flanked by the 5’ and 3’ regulatory sequences of *Mla6*. The two binary vectors pANIC12A-*Scs6* and pBract202-pMla6-*Scs6*-tMla6 were introduced into barley cv. Golden Promise and DH47 by *Agrobacterium-mediated* transformation following the methods described by (23, 24), respectively.

### Transient gene expression in *N. benthamiana* and protein detection by immunoblotting

Generation of entry and destination vectors of MLA1, MLA6, MLA22, and AVR_A1_ and AVR_A6_ is described in (25, 26). The wild-type coding sequence without the stop codon of *Scs6* and of *MLA16*, *MLA18*-1 and *MLA25* (27) was amplified by PCR using attB-primers followed by BP reaction into pDONR221 to generate a gateway-compatible entry clone (**Table S6**). Entry vectors carrying wild-type cDNAs of *MLA3*, *FT153*, *FT352-2* and *MLA18-2* without stop codons and insect-cell codon-optimized *Sr50* were obtained by gene synthesis (GeneArt, Invitrogen). Plasmids encoding chimeric SCS6/MLA1 and SCS6/MLA6 receptors were assembled using the NEBuilder HiFi assembly Kit (NEB) based on the domain boundaries reported in (28). pENTR221-*Scs6* was used as a template to generate *Scs6*^S793F^ and *Scs6*^H510V^ via PCR mutagenesis using the Q5 Site-Directed Mutagenesis Kit (NEB).

LR-Clonase II (Thermo Fisher) was used to recombine the genes into the expression vector pGWB517 that carries a C-terminal linker region followed by an in-frame 4xmyc epitope tag (29). The integrity of all entry and destination vectors was confirmed by whole-plasmid nanopore sequencing (Eurofins). Expression constructs were transformed into *Agrobacterium tumefaciens* GV3101 (pMP90RK) by electroporation. Transformants were selected for three days at 28 °C on LB agar medium containing rifampicin (15 mg ml^-1^), gentamycin (25 mg ml^-1^), kanamycin (50 mg ml^-1^), and spectinomycin (50 mg ml^-1^). Transformants were cultured in liquid LB medium containing the corresponding antibiotics at 28 h overnight, after which they were harvested by centrifugation at 2500 g for 6 minutes and resuspended in infiltration buffer (10 mM MES pH 5.6, 10 mM MgCl_2_, and 200 μM acetosyringone). Transient gene expression in leaves of four-week-old *N. benthamiana* plants was performed via Agrobacterium-mediated transient expression assays in the presence of the P19 and CMV2b suppressors of RNAi silencing (30). The final OD_600_ of bacteria carrying expression vectors of immune receptors and silencing suppressors was set to 0.5, unless stated otherwise. For the expression of effector proteins, the OD_600_ was increased to 1.0 unless stated differently. Twenty-four hours after *agrobacterium*-mediated gene delivery, IWF was infiltrated, as indicated. For this, a small subset towards the outer part of the region of transgene expression was infiltrated with approx. 25–50 μL of IWF. Cell death phenotypes were assessed and documented at 2 or 5 days after agroinfiltration for IWF-triggered cell death or effector-triggered cell death, respectively.

For the detection of protein accumulation, leaf material of four individual plants was harvested 48 h after infiltration, flash-frozen in liquid nitrogen and ground to powder using a Retsch bead beater. Then, 100 mg plant tissue powder was resuspended in 200 μL Urea-SDS sample buffer (50 mM Tris-HCl pH 6.8, 2% SDS, 8 M Urea, 4% β-mercaptoethanol, 5% Glycerol and 0.004% bromophenol blue) and vortexed at room temperature for 10 min. After centrifugation at 16,000 g for 15 min, 10 μl of supernatant were loaded onto a 10% SDS-PAGE without prior boiling. Separated proteins were transferred to a PVDF membrane and probed with monoclonal mouse anti-Myc (1:3,000; R950-25, Thermofisher), polyclonal rabbit anti-GFP (1:3,000; pabg1, Chromotek) followed by polyclonal goat anti-mouse IgG-HRP (1:7,500; ab6728, Abcam) or polyclonal swine anti-rabbit IgG-HRP (1:5,000; PO399, Agilent DAKO) antibodies. Myc-tagged proteins were detected using SuperSignal West Femto: SuperSignal substrates (ThermoFisher Scientific) in a 1:1 ratio. SuperSignal Femto Substrate was used for AVR_A1_ and SuperSignal Substrate for AVR_A6_.

### Sequencing of *Scs6* homologs in cultivated and wild barley accessions

The primer pair SCS6-F2 and SCS6-R17 (**Table S6**) was used to amplify the whole gene of *Scs6* in cultivated and wild barley accessions (**Table S4**). PCR products were purified using Quick PCR Purification Kit (Invitrogen, Carlsbad, CA) and sequenced by EurofinGenomics (Louisville, KY) using primers F2, R2, R3, SCS6-Seq-R1, SCS6-Seq-F1, and SCS6-Seq-F2 (**Table S6**). Homologs were aligned against the CDS of *Scs6* and any single nucleotide polymorphism (SNP) was validated by checking the sequence quality manually. Finally, the sequences of *Scs6* homologs excluding introns were translated into amino acid sequences and used for phylogenetic analysis.

### Phylogenetic analysis of *Scs6* and *Mla* alleles

Previously published MLA protein sequences were retrieved from NCBI and aligned via SnapGene using Clustal Omega. Protein sequences of SCS6 variants in wild barley identified in this study were manually added to the alignment (**Supplementary Data S1 and S3**). A BLAST search was conducted to identify MLA-like sequences in the Triticeae using MLA1 and SCS6 as an input. The identified candidate sequences were manually inspected to remove truncated (> 840 aa) sequences. The resulting alignment was used to generate neighbor-net networks as described in (31) using splitstree4 (32). For the phylogenetic analyses of individual SCS6/MLA domains, we regarded the first N-terminal 161 amino acids that align with SCS6 as the CC domain, the sequence stretching from amino acid 162 to 551 as the NB domain, and the sequence from amino acid 551 to the end as the LRR. To analyze sites undergoing positive selection, the Clustal alignment of protein sequences as well as the corresponding nucleotide coding sequences were used as an input for PAL2NAL to generate a codon-aware MSA **(Supplementary Data S1**). In this MSA, sites under episodic positive selection were identified using the MEME algorithm (33) with default parameters and sites under pervasive positive selection identified using FUBAR (34) with default settings. Both MEME as well as FUBAR were accessed via the datamonkey application (35).

### Geographic distribution of wild barley accessions susceptible to *Bs*_ND90Pr_

The geographic coordinates of sampled accessions from the WBDC (5) and (31) were plotted in QGIS 3.32. Geographic vector map datasets were downloaded from the Natural Earth repository (http://www.naturalearthdata.com).

## Acknowledgements

The authors thank Joseph Mullins for assistance in disease phenotyping experiments in greenhouse, Inmaculada Hernandez-Pinzon and Xiaohong Jiang for barley transformation, Megan Overlander-Chen for taking care of transgenic barley plants, Antonín Dreiseitl for testing Bowman for reaction to barley powdery mildew isolates, Maria von Korff for providing some of the wild barley accessions used in the study. We also thank P. Cápal, M. Said, Z. Dubská, J. Weiserová, and E. Jahnová for assistance with chromosome flow sorting and preparation of chromosome DNA. This research was funded by the Triticeae-CAP project (2011-68002-30029) of the US Department of Agriculture National Institute of Food and Agriculture (S.Z.), the Marie Curie Fellowship grant ‘AEGILWHEAT’ (H2020-MSCA-IF-2016-746253) (I.M), the Hungarian National Research, Development and Innovation Office (K135057) (I.M.), the Max-Planck-Gesellschaft (P.S.-L.), the Deutsche Forschungsgemeinschaft (DFG, German Research Foundation) in the Collaborative Research Centre Grant (SFB-1403 – 414786233 B08) (P.S.-L.), Germany’s Excellence Strategy CEPLAS (EXC-2048/1, project 390686111) (P.S.-L.), the Gatsby Charitable Foundation (M.J.M.), and the United States Department of Agriculture-Agricultural Research Service CRIS #5062-21220-025-000D (M.J.M.) and 3060-21000-046-000D (S.Y.). Mention of trade names or commercial products in this publication is solely for the purpose of providing specific information and does not imply recommendation or endorsement by the U.S. Department of Agriculture. USDA is an equal opportunity provider and employer.

## Author Contributions

Y.L., F.K., P.S.-L. and S.Z. designed research; Y.L., F.K., M.Z., I.M., E.L., P.K., P.X., S.Y., M.J.M., S.M. performed research; J.D., J.D.F., Y.D., B.S., S.M., B.J.S contributed new reagents/resources/analytic tools; Y.L., F.K., S.Y., P.S.-L. and S.Z. analyzed data; and F.K., Y.L., P.S.-L. and S.Z. wrote the paper. All reviewed the manuscript.

---

**Figure S1.**
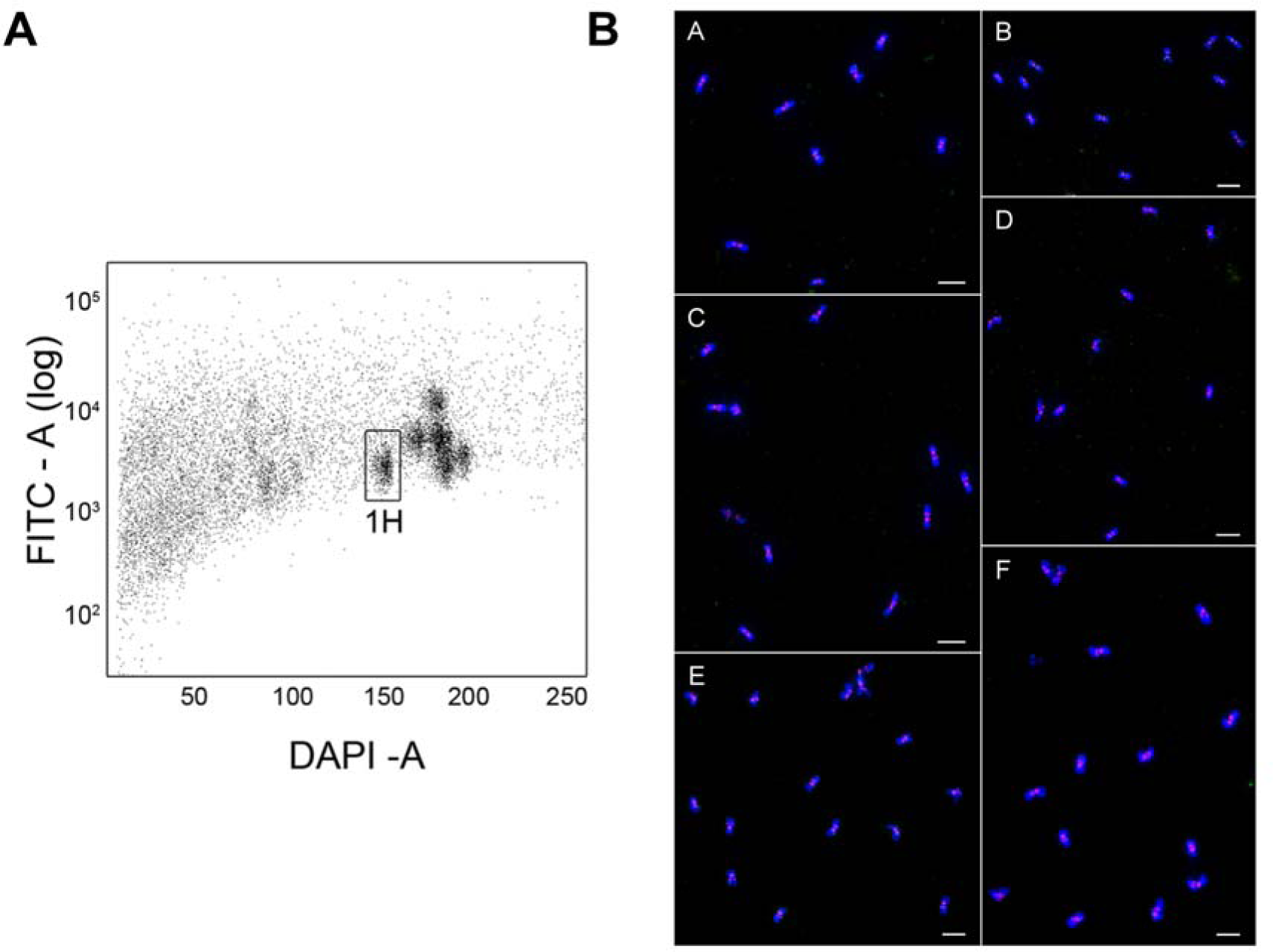
Chromosome flow sorting of 1H. (A) Bivariate flow karyotype of mitotic metaphase chromosomes isolated from barley cv. Bowman. DAPI-A vs. FITC-A dot plot was obtained after the analysis of DAPI-stained chromosome suspensions labeled by FISHIS with FITC-conjugated probes for GAA microsatellites. 1H chromosomes were sorted from the sort window shown as rectangle at purities of 87-96%. (B), Chromosome 1H flow-sorted from barley cv. Bowman a) and its mutants EMS14 b), EMS494 c), EMS621 d), EMS1317 e) and EMS1473 f). The chromosomes were flow-sorted onto microscope slides, counterstained by DAPI (blue) and identified by fluorescence *in situ* hybridization with s probe GAA microsatellite repeats (red). Bar = 10 mm.

**Figure S2.**
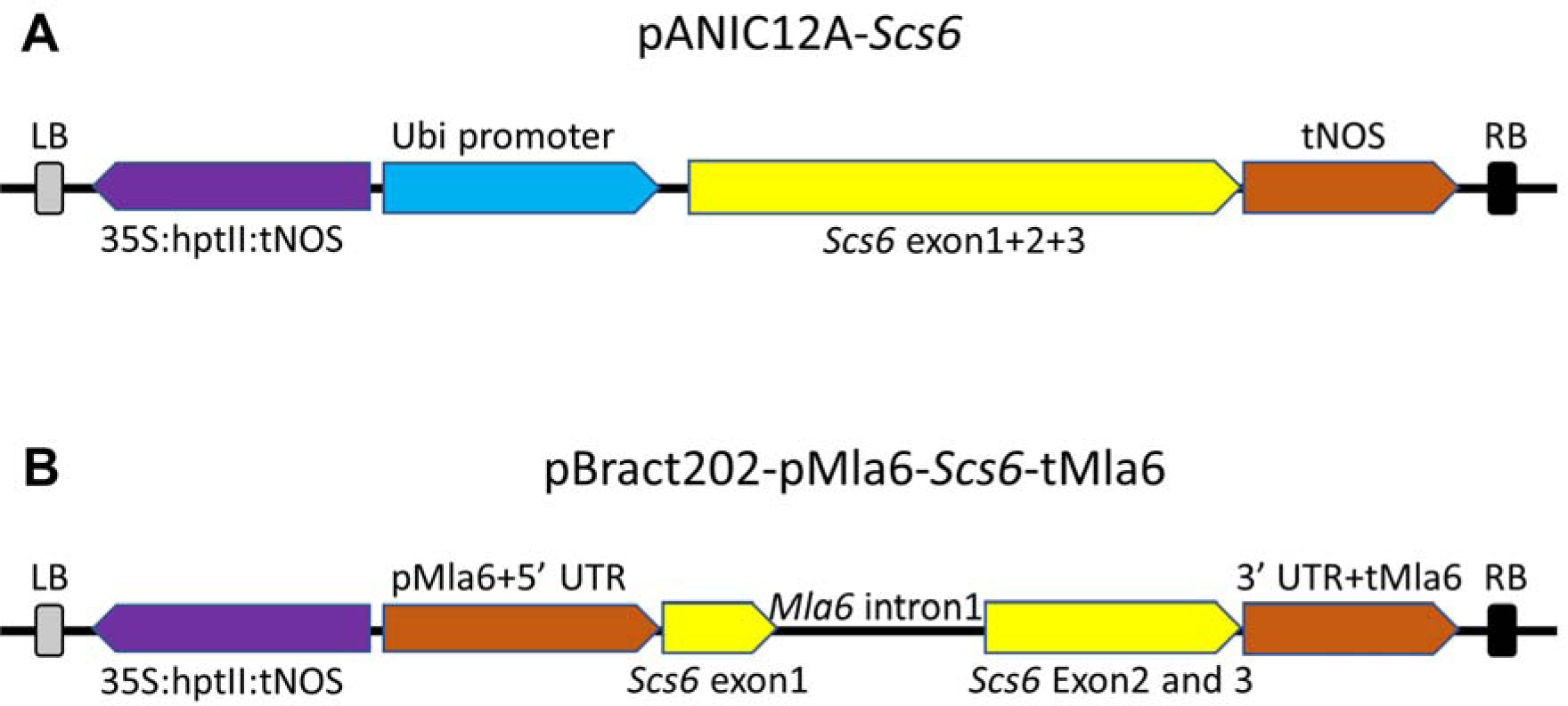
Gene constructs used for transformation of barley cv. Golden Promise and barley line SxGP DH-47. (A) The coding sequence of *Scs6* was expressed under the maize Ubi promoter and NOS terminator (NOSt), and cloned into the backbone of binary vector pANIC12A (21). (B) The coding sequence of *Scs6* was expressed under the promoter+5’UTR (pMla6+5’ UTR) and 3’UTR+terminator (3’ UTR+tMla6) of *Mla6*, which was assembled into binary vector pBract202 (22). Both constructs use hptII driven by the 35S Cauliflower Mosaic Virus (35S) promoter for plant selection during transformation (shown in purple). Left and right T-DNA borders (LB and RB) are shown with filled grey and black blocks, respectively.

**Figure S3.**
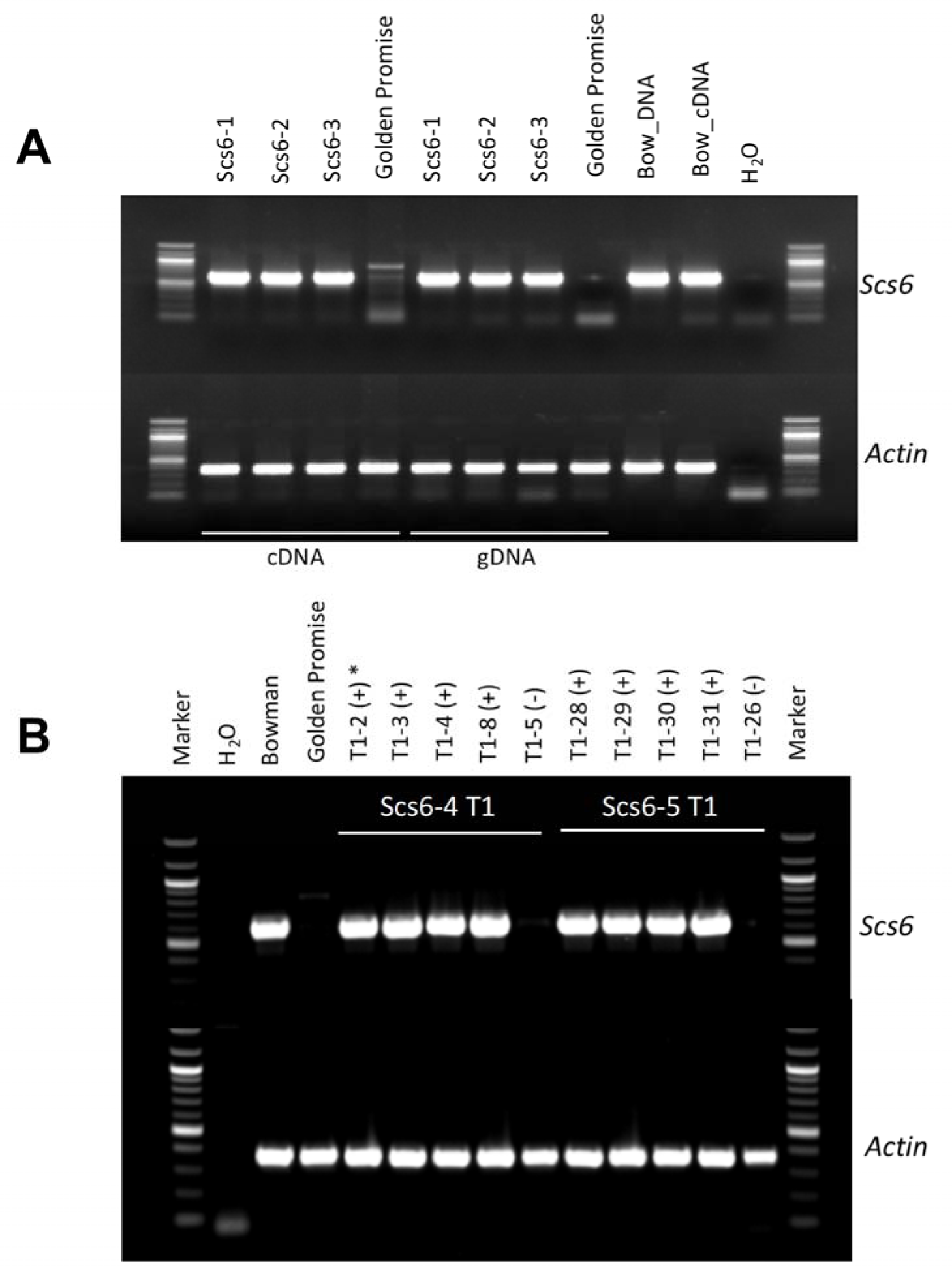
PCR analysis of *Scs6* transgenic barley plants. (A) PCR amplification of *Scs6* with genomic DNA and cDNA from Bowman (Bow_DNA and Bow_cDNA), Golden Promise (GP_DNA and GP_cDNA) and three T0 transgenic barley plants (GP_Scs6-1, GP_Scs6-2, and GP_Scs6-3) using primers Scs6-F4+Scs6-R1 (Table S6). (B) PCR amplification of *Scs6* with cDNA derived from Bowman (Bow_cDNA), Golden Promise (GP_cDNA), and five T1 plants derived from each of the two T0 transgenic barley plants (GP_Scs6-4 and GP_Scs6-5) using primers Scs6-F4+Scs6-R1 (Table S6). * (+) indicated that PCR was positive and (-) indicated that PCR was negative.

**Figure S4.**
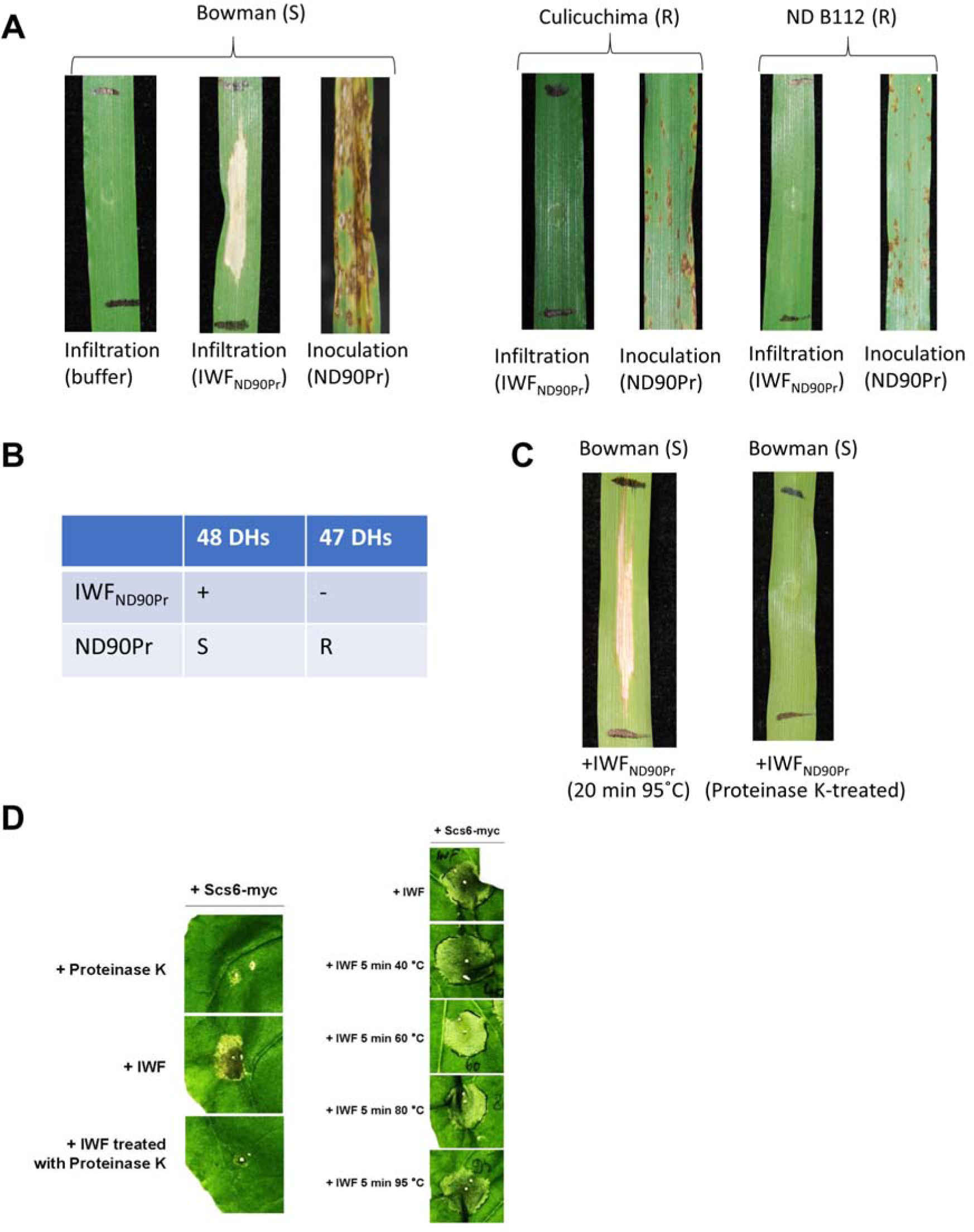
Partial characterization of the *Bipolaris sorokiniana* isolate ND90Pr NPS1-derived effector. (A) Reactions of different barley genotypes to *B. sorokiniana* isolate ND90Pr at seven days after inoculation or infiltration of intracellular washing fluid (IWF_ND90Pr_) extracted from ND90Pr-infected Bowman leaves. B. Sensitivity (‘+’) to IWF_ND90Pr_ is coupled with susceptibility (‘S’) to ND90Pr inoculation in the doubled haploid (DH) mapping population derived from the cross between Bowman-BC and Culicuchima. (C-D) The NPS1-derived effector is heat-resistant but sensitive to proteinase K-treatment.

**Figure S5.**
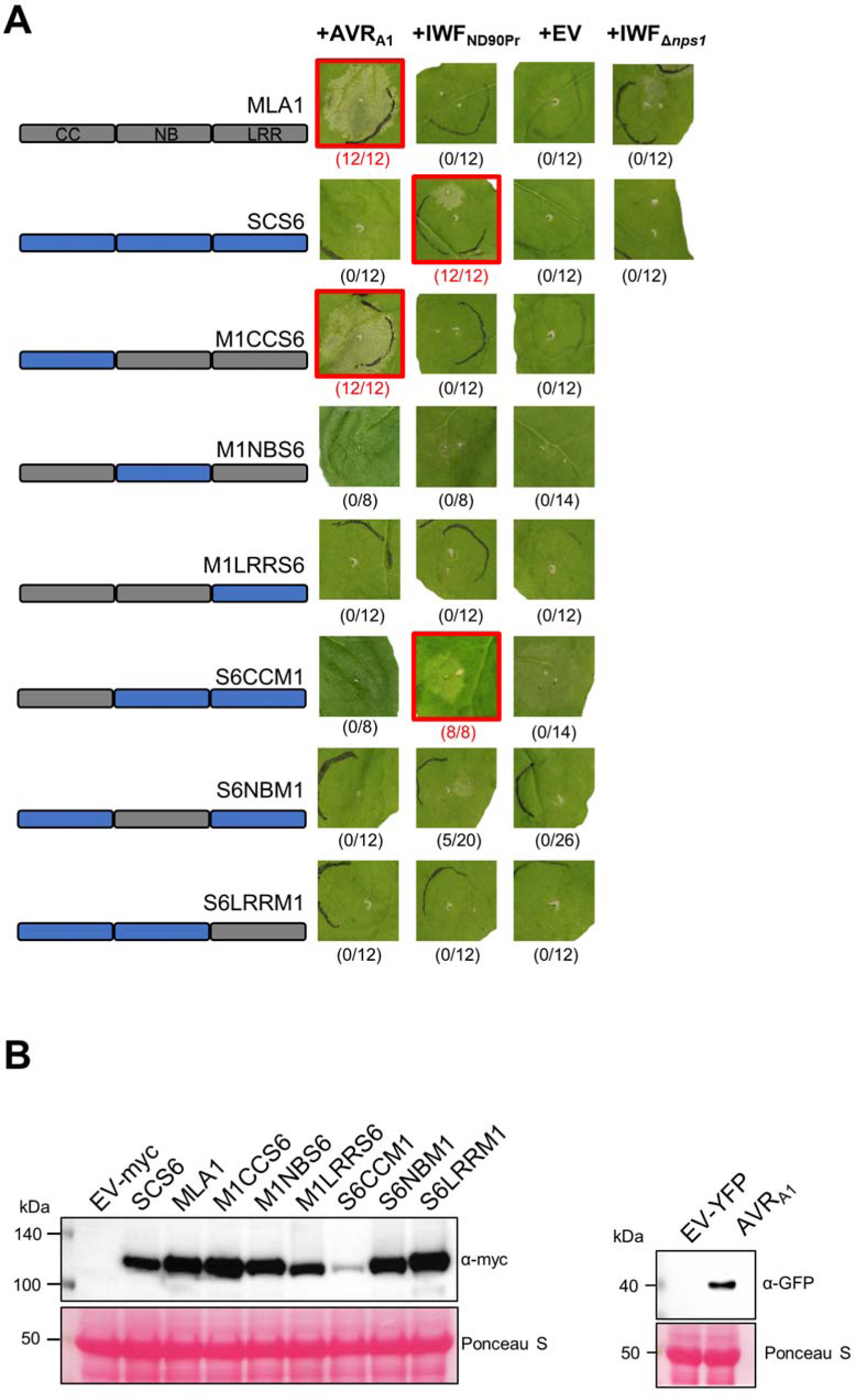
A *Bipolaris sorokiniana* ND90Pr effector activates SCS6 to cause cell death in *Nicotiana benthamiana* depending on its NB and LRR domain. (A) *Nicotiana benthamiana* plants were transformed transiently, as indicated. AVR_A1_ does not carry a signal peptide. Twenty-four hours after *Agrobacterium*-mediated gene delivery, IWF_ND90Pr_ or IWF_Δ*nps1*_ was infiltrated, as indicated. Cell death phenotypes were assessed and documented at two or four days after agroinfiltration for IWF-triggered cell death or effector-triggered cell death, respectively. Representative pictures of at least eight biological replicates (indicated in brackets) are shown and combinations that resulted in cell death are highlighted with a red box. All gene constructs were transformed by setting the OD_600_ of *A. tumefaciens* to 0.5, except S6CCM1, which was reduced to 0.2 to attenuate auto-activity. (B). Protein levels of receptor-4xMyc (approx. 114 kDa and AVR_A1_-mYFP (39 kDA) in *Nicotiana benthamiana*.

**Figure S6.**
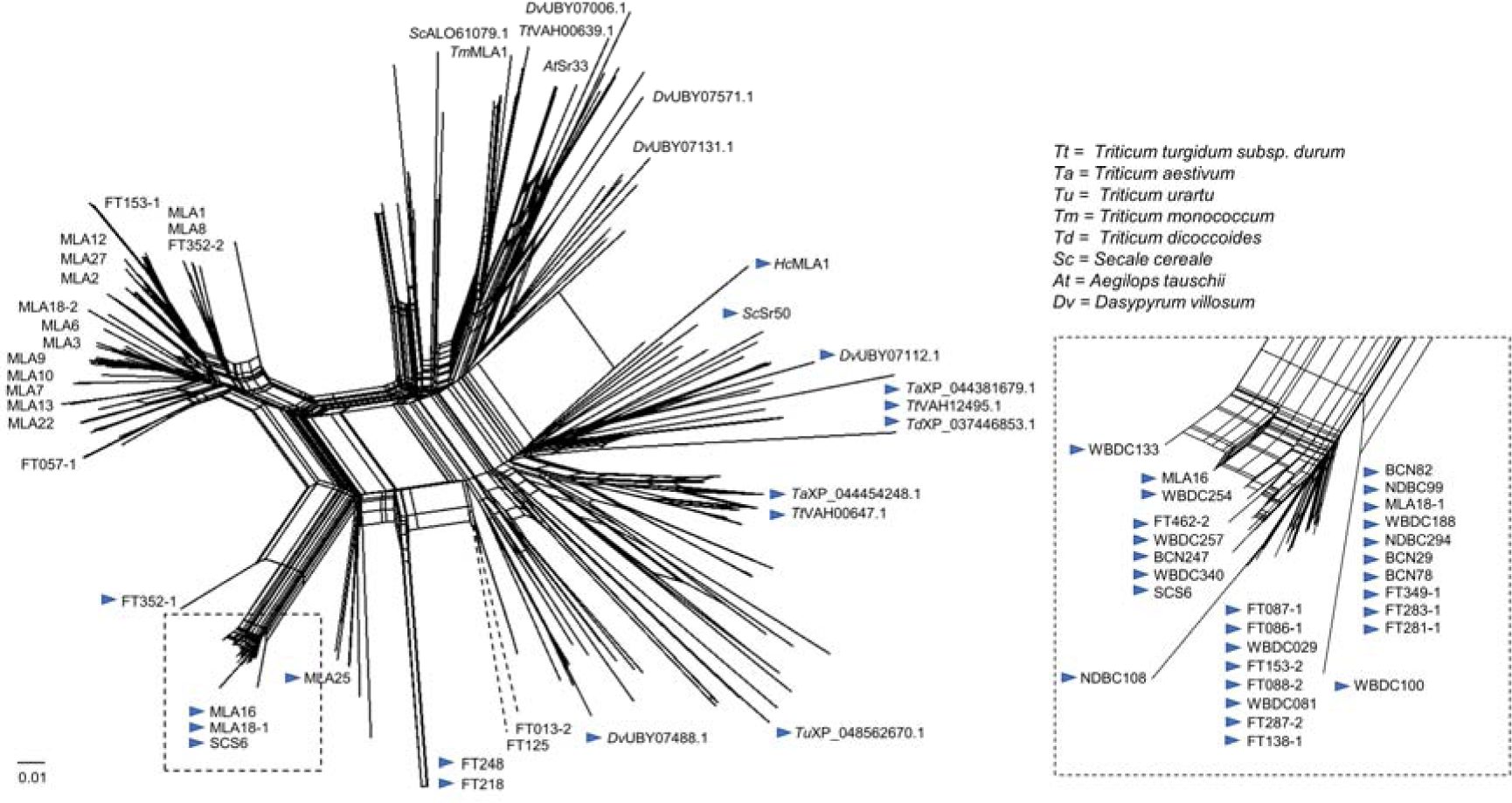
Phylogenetic tree including RGH1 sequences identified in members of the Triticeae. Neighbor-Net analysis of 207 RGH1 protein sequences including 28 previously identified MLA proteins from barley (27), 50 sequences from wild barley (31), as well as additional sequences from wild or domesticated barley identified in this this study. Sequences identified in *Dasypyrum villosum* are based on (36). Additional RGH1 sequences outside the genus *Hordeum* were identified using BLASTp. MLA subfamily 2 members are highlighted using a blue triangle. Dashed lines indicate that the respective branch was reduced to 50% of its size.

**Figure S7.**
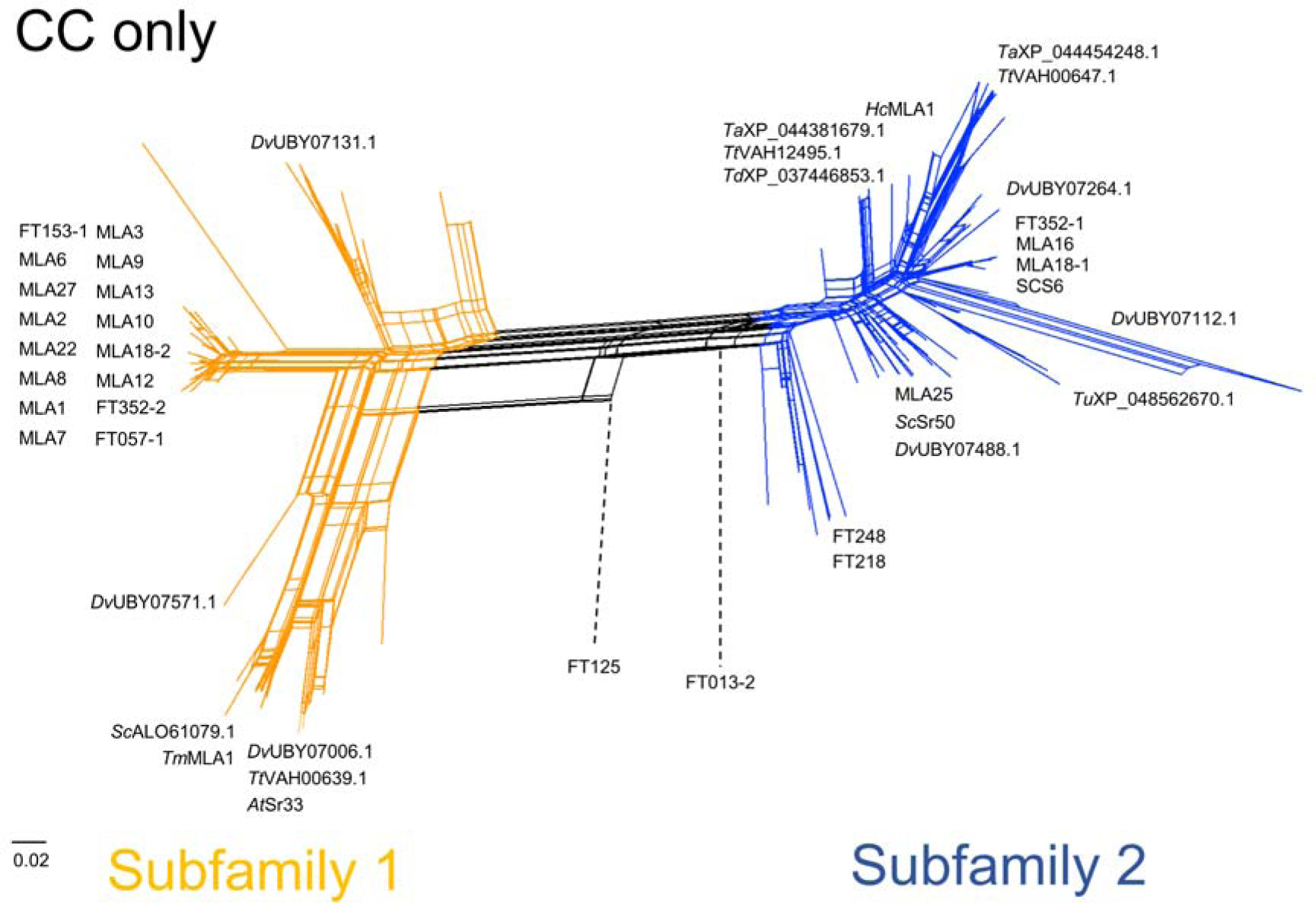
Phylogenetic tree of CC domains of RGH1 members identified in members of the Triticeae. Neighbor-Net analysis of 207 RGH1 protein sequences including 28 previously identified MLA proteins from barley (27), 50 sequences from wild barley (31), as well as additional sequences from wild or domesticated barley identified in this this study. Sequences identified in *Dasypyrum villosum* are based on (36). Bootsstrap support for the separation of the two Subfamilies: 100% (1000 iterations). Dashed lines indicate that the respective branch was reduced to 50% of its size.

**Figure S8.**
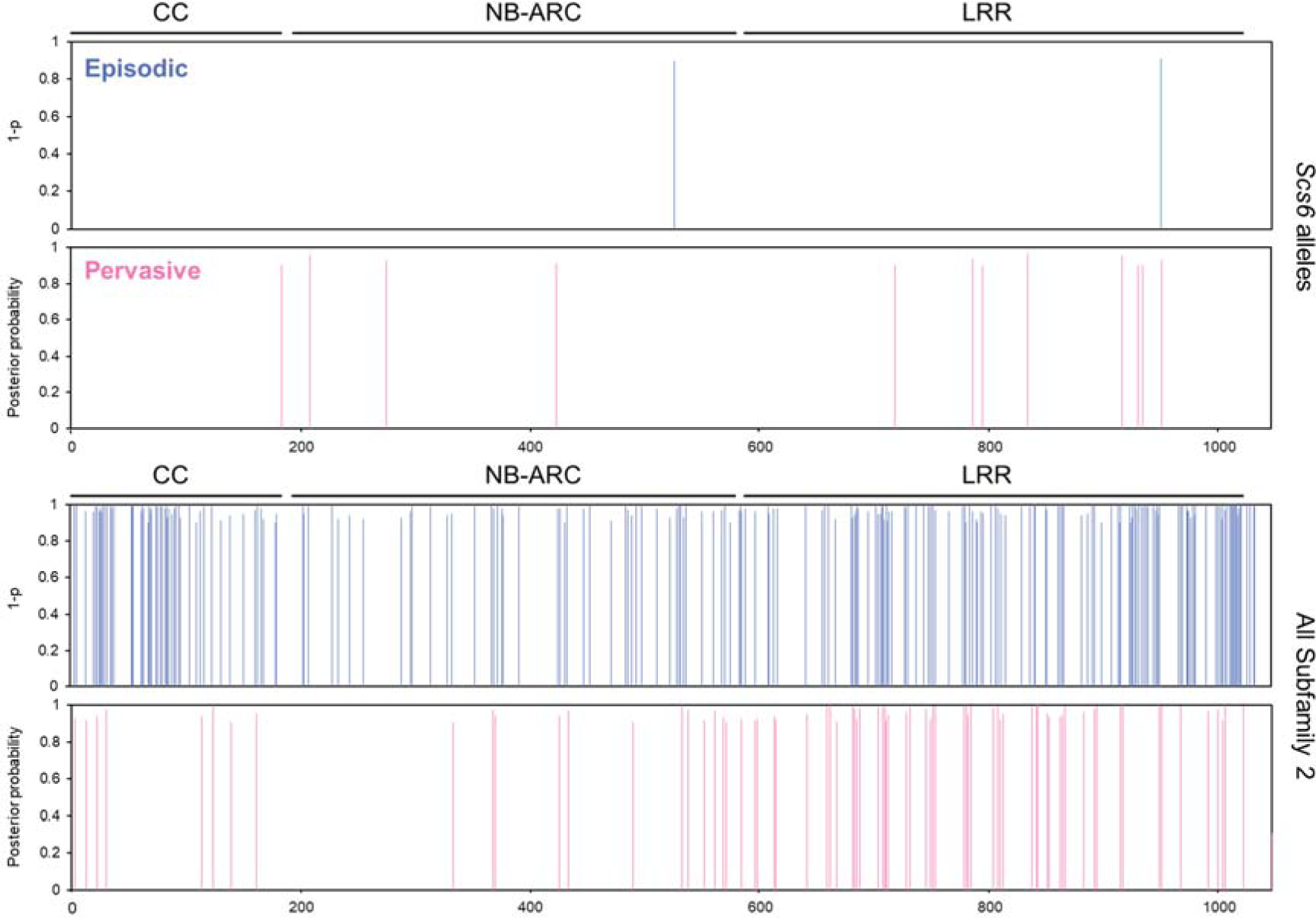
Identification of sites under positive selection in SCS6 and all MLA subfamily 2 members from the Triticeae. Sites under episodic (blue bars) or pervasive (pink bars) positive selection were identified using MEME or FUBAR, respectively. Analysis was performed as described in (31). For the analysis of MLA Subfamily 2 members, 112 sequences that were identified via BLASTp and belonging to MLA Subfamily 2 based on Figure S7 were included in the analysis.

**Table S1.** Summary of chromosome 1H sorting and sequencing of wild type and EMS mutants.

| Sample | Purity of Chromosome 1H | Amount of DNA after MDA (ug) | Number of reads <sup>a</sup> | Coverage of Chromosome 1H <sup>b</sup> |
| --- | --- | --- | --- | --- |
| Bowman | 87.70% | 5.56 | 338,824,744 | 61 |
| EMS14 | 90.90% | 1.94 | 439,682,584 | 79 |
| EMS494 | 88.50% | 5.38 | 470,198,020 | 84 |
| EMS621 | 96.70% | 2.14 | 469,926,034 | 84 |
| EMS1317 | 93.45% | 5.45 | 470,243,174 | 84 |
| EMS1473 | 96.07% | 5.58 | 453,842,418 | 81 |

<sup>a</sup> The total number of clean reads of 100 base pair.
<sup>b</sup> The coverage was calculated based on the length of chromosome 1H of reference genome Morex (558,535,432 bp).

**Table S2.** Mutation overlap in contigs from flow-sorted 1H chromosome of barley.

| Barley 1H<sup>a</sup> | Number of contigs |
| --- | --- |
| Mutated in 0 line | 126353 |
| Mutated in 1 line | 2842 |
| Mutated in 2 lines | 87 |
| Mutated in 3 lines | 4 |
| Mutated in 4 lines | 1 |
| Mutated in 5 lines | 0 |

<sup>a</sup> The number of total assembled contigs of chromosome 1H of wild type.

**Table S3.** Copy number of the transgene in T1 progeny derived from barley line SxGP DH47 and their segregating infection responses to *Bipolaris soroiniana* isolate ND90Pr.

| ID | CallusID | Generation | T-DNA Insert* | R : IM : S ** |
| --- | --- | --- | --- | --- |
| HVT_04655 | 1-1 | T1 | 3 copies | 11 : 0 : 46 |
| HVT_04656 | 1-2 | T1 | 3 copies | 33 : 0 : 92 |
| HVT_04657 | 1-3 | T1 | 3 copies | 16 : 0 : 53 |
| HVT_04658 | 1-4 | T1 | 3 copies | 27 : 0 : 97 |
| HVT_04659 | 2-1 | T1 | 1 copy | 28 : 23 : 36 |
| HVT_04660 | 2-2 | T1 | 1 copy | 33 : 17 : 12 |
| HVT_04682 | 3-1 | T1 | 2 copies | 10 : 3 : 34 |
| HVT_04683 | 4-1 | T1 | 5 copies | 3 : 1 : 5 |
| HVT_04684 | 4-2 | T1 | 5 copies | 13 : 0 : 21 |
| HVT_04686 | 5-2 | T1 | 2 copies | 25 : 36 : 33 |
| HVT_04688 | 7-1 | T1 | 2 copies | 0 : 0 : 23 |
| HVT_04689 | 7-2 | T1 | 2 copies | 0 : 14 : 11 |
| HVT_04690 | 8-1 | T1 | 1 copy | 16 : 5 : 45 |
| HVT_04692 | 10-1 | T1 | 3 copies | 0 : 1 : 19 |
| HVT_04693 | 11-1 | T1 | 1 copy | 21 : 2 : 45 |
| HVT_04694 | 11-2 | T1 | 1 copy | 10 : 0 : 23 |
| HVT_04695 | 11-3 | T1 | 2 copies | 7 : 0 : 29 |
| HVT_04696 | 12-1 | T1 | 2 copies | 28 : 18 : 69 |
| HVT_04697 | 12-2 | T1 | 5 copies | 2 : 1 : 77 |

\* Determined by qPCR for the Hygromycin resistant gene
\*\* Infection response of T1 plants against isolate ND90Pr of *B. sorokiniana*. R indicates resistance with disease rating 1-3; IM indicates an intermediate reaction with disease rating 4-6; S indicates susceptibility with disease rating 7-9.

**Table S4.**
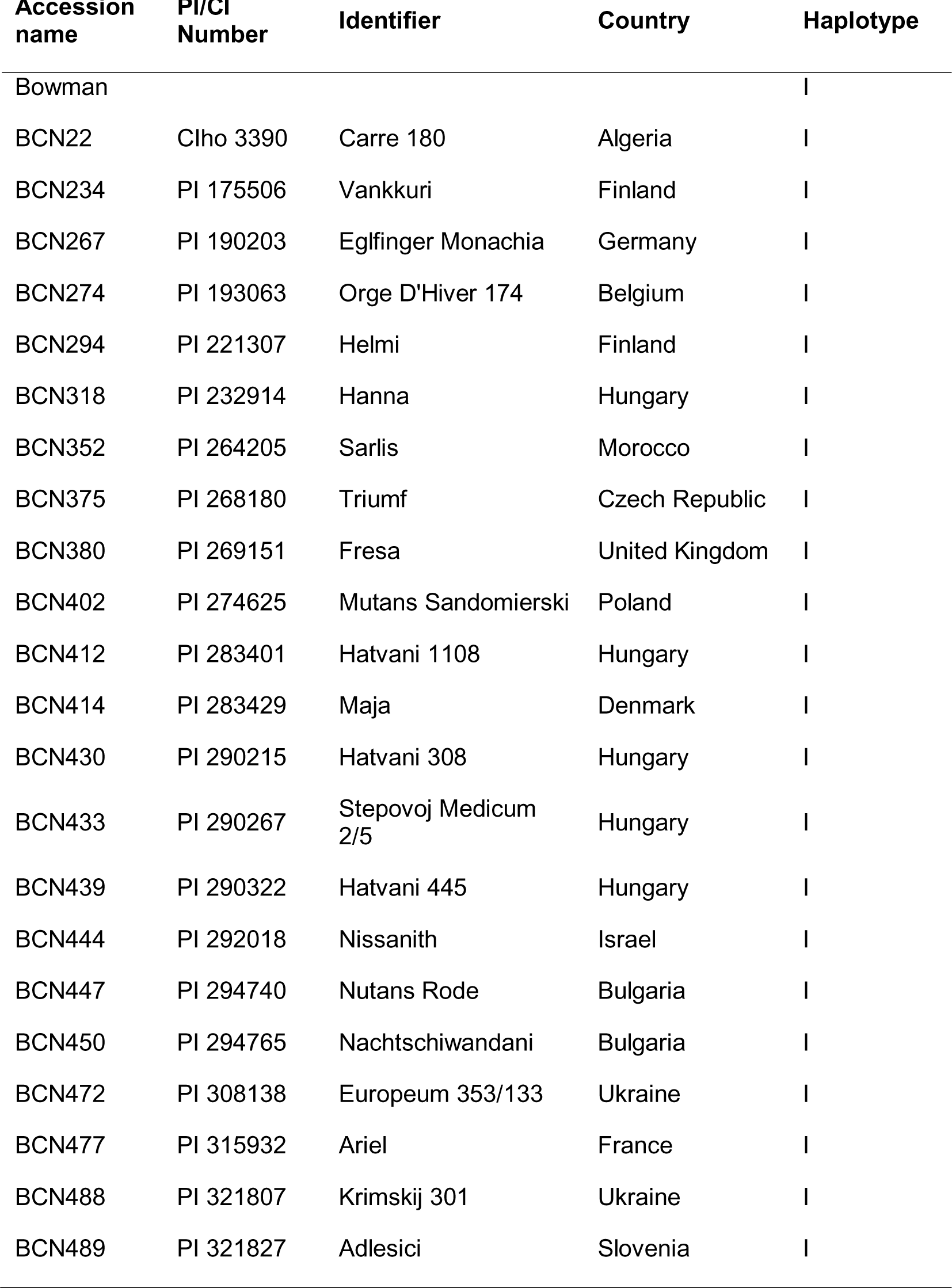
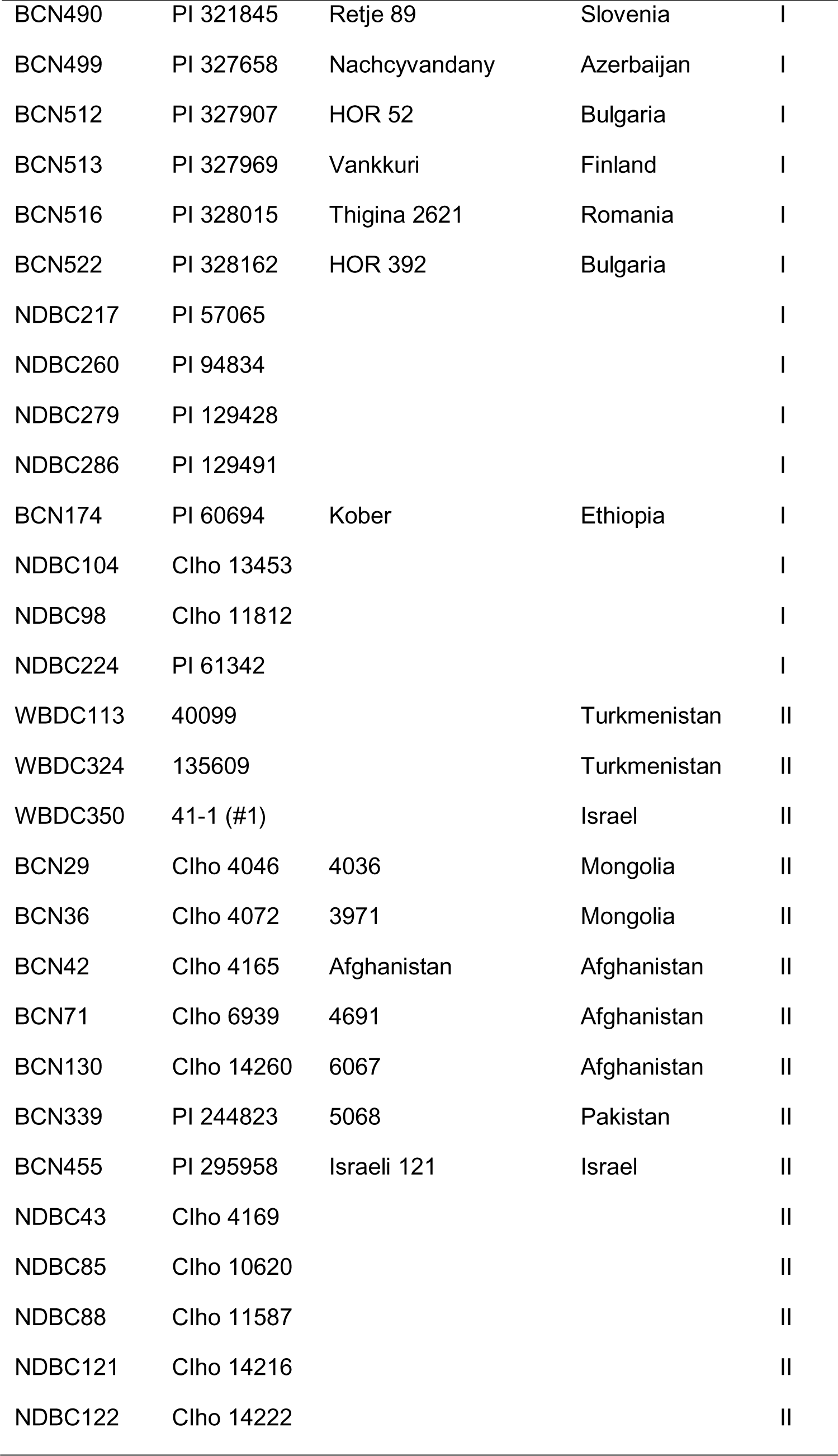
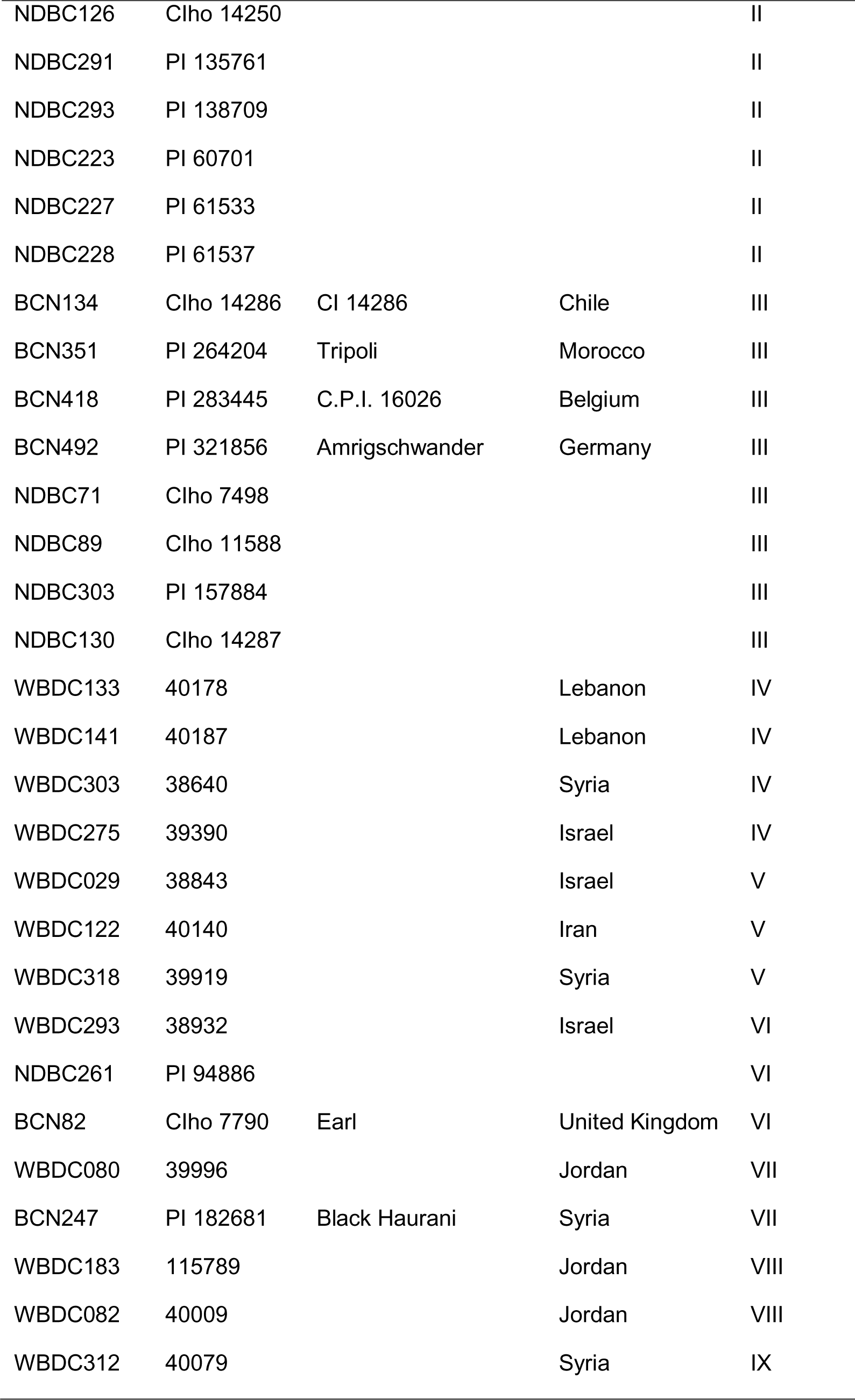
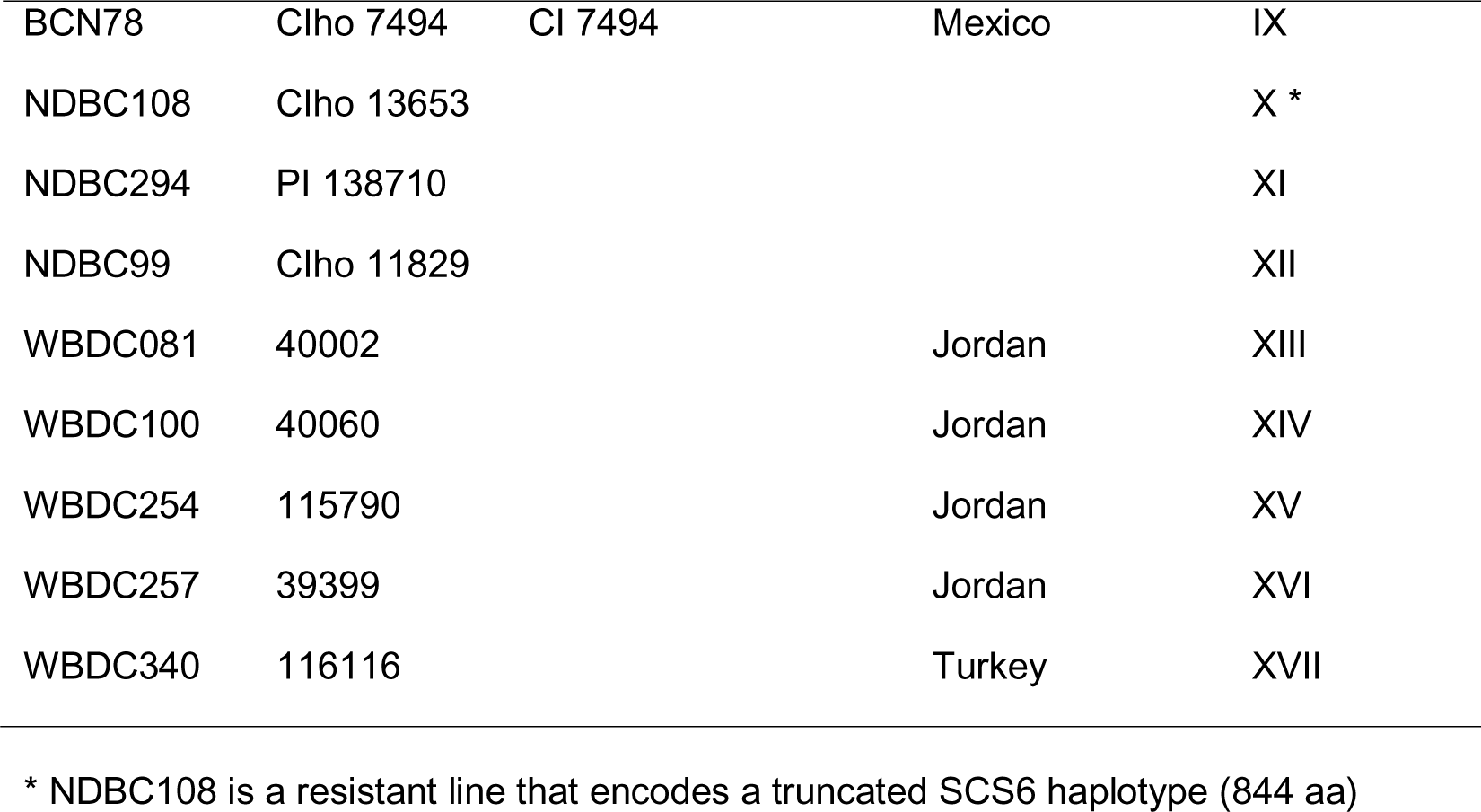
Fully sequenced *Scs6* haplotypes found in wild and domesticated barley lines.

| Accession name | PI/CI Number | Identifier | Country | Haplotype |
| --- | --- | --- | --- | --- |
| Bowman |  |  |  | I |
| BCN22 | CIho 3390 | Carre 180 | Algeria | I |
| BCN234 | PI 175506 | Vankkuri | Finland | I |
| BCN267 | PI 190203 | Eglfinger Monachia | Germany | I |
| BCN274 | PI 193063 | Orge D'Hiver 174 | Belgium | I |
| BCN294 | PI 221307 | Helmi | Finland | I |
| BCN318 | PI 232914 | Hanna | Hungary | I |
| BCN352 | PI 264205 | Sarlis | Morocco | I |
| BCN375 | PI 268180 | Triumf | Czech Republic | I |
| BCN380 | PI 269151 | Fresa | United Kingdom | I |
| BCN402 | PI 274625 | Mutans Sandomierski | Poland | I |
| BCN412 | PI 283401 | Hatvani 1108 | Hungary | I |
| BCN414 | PI 283429 | Maja | Denmark | I |
| BCN430 | PI 290215 | Hatvani 308 | Hungary | I |
| BCN433 | PI 290267 | Stepovoj Medicum 2/5 | Hungary | I |
| BCN439 | PI 290322 | Hatvani 445 | Hungary | I |
| BCN444 | PI 292018 | Nissanith | Israel | I |
| BCN447 | PI 294740 | Nutans Rode | Bulgaria | I |
| BCN450 | PI 294765 | Nachtschiwandani | Bulgaria | I |
| BCN472 | PI 308138 | Europeum 353/133 | Ukraine | I |
| BCN477 | PI 315932 | Ariel | France | I |
| BCN488 | PI 321807 | Krimskij 301 | Ukraine | I |
| BCN489 | PI 321827 | Adlesici | Slovenia | I |
| BCN490 | PI 321845 | Retje 89 | Slovenia | I |
| BCN499 | PI 327658 | Nachcyvandany | Azerbaijan | I |
| BCN512 | PI 327907 | HOR 52 | Bulgaria | I |
| BCN513 | PI 327969 | Vankkuri | Finland | I |
| BCN516 | PI 328015 | Thigina 2621 | Romania | I |
| BCN522 | PI 328162 | HOR 392 | Bulgaria | I |
| NDBC217 | PI 57065 |  |  | I |
| NDBC260 | PI 94834 |  |  | I |
| NDBC279 | PI 129428 |  |  | I |
| NDBC286 | PI 129491 |  |  | I |
| BCN174 | PI 60694 | Kober | Ethiopia | I |
| NDBC104 | Clho 13453 |  |  | I |
| NDBC98 | Clho 11812 |  |  | I |
| NDBC224 | PI 61342 |  |  | I |
| WBDC113 | 40099 |  | Turkmenistan | II |
| WBDC324 | 135609 |  | Turkmenistan | II |
| WBDC350 | 41-1 (#1) |  | Israel | II |
| BCN29 | Clho 4046 | 4036 | Mongolia | II |
| BCN36 | Clho 4072 | 3971 | Mongolia | II |
| BCN42 | Clho 4165 | Afghanistan | Afghanistan | II |
| BCN71 | Clho 6939 | 4691 | Afghanistan | II |
| BCN130 | Clho 14260 | 6067 | Afghanistan | II |
| BCN339 | PI 244823 | 5068 | Pakistan | II |
| BCN455 | PI 295958 | Israeli 121 | Israel | II |
| NDBC43 | Clho 4169 |  |  | II |
| NDBC85 | Clho 10620 |  |  | II |
| NDBC88 | Clho 11587 |  |  | II |
| NDBC121 | Clho 14216 |  |  | II |
| NDBC122 | Clho 14222 |  |  | II |
| NDBC126 | Clho 14250 |  |  | II |
| NDBC291 | PI 135761 |  |  | II |
| NDBC293 | PI 138709 |  |  | II |
| NDBC223 | PI 60701 |  |  | II |
| NDBC227 | PI 61533 |  |  | II |
| NDBC228 | PI 61537 |  |  | II |
| BCN134 | Clho 14286 | CI 14286 | Chile | III |
| BCN351 | PI 264204 | Tripoli | Morocco | III |
| BCN418 | PI 283445 | C.P.I. 16026 | Belgium | III |
| BCN492 | PI 321856 | Amrigschwander | Germany | III |
| NDBC71 | Clho 7498 |  |  | III |
| NDBC89 | Clho 11588 |  |  | III |
| NDBC303 | PI 157884 |  |  | III |
| NDBC130 | Clho 14287 |  |  | III |
| WBDC133 | 40178 |  | Lebanon | IV |
| WBDC141 | 40187 |  | Lebanon | IV |
| WBDC303 | 38640 |  | Syria | IV |
| WBDC275 | 39390 |  | Israel | IV |
| WBDC029 | 38843 |  | Israel | V |
| WBDC122 | 40140 |  | Iran | V |
| WBDC318 | 39919 |  | Syria | V |
| WBDC293 | 38932 |  | Israel | VI |
| NDBC261 | PI 94886 |  |  | VI |
| BCN82 | Clho 7790 | Earl | United Kingdom | VI |
| WBDC080 | 39996 |  | Jordan | VII |
| BCN247 | PI 182681 | Black Haurani | Syria | VII |
| WBDC183 | 115789 |  | Jordan | VIII |
| WBDC082 | 40009 |  | Jordan | VIII |
| WBDC312 | 40079 |  | Syria | IX |
| BCN78 | Clho 7494 | CI 7494 | Mexico | IX |
| NDBC108 | Clho 13653 |  |  | X * |
| NDBC294 | PI 138710 |  |  | XI |
| NDBC99 | Clho 11829 |  |  | XII |
| WBDC081 | 40002 |  | Jordan | XIII |
| WBDC100 | 40060 |  | Jordan | XIV |
| WBDC254 | 115790 |  | Jordan | XV |
| WBDC257 | 39399 |  | Jordan | XVI |
| WBDC340 | 116116 |  | Turkey | XVII |

\* NDBC108 is a resistant line that encodes a truncated SCS6 haplotype (844 aa)

**Table S5.** Summary of 50 wild barley lines used in previous study and their reaction to *Bipolaris sorokiniana* ND90Pr. Modified based on 1).

| Sample | Line_ID | Origin<sup>a</sup> | Latitude | Longitude | Population | No. of *Rgh1/Mla* | Subfamily 1 | Subfamily 2 | Other | Reaction to *B. sorokiniana*<sup>c</sup> |
| --- | --- | --- | --- | --- | --- | --- | --- | --- | --- | --- |
| FT010 | B1K-04-08 | ISR | 31.834371 | 35.305292 | JJ | – | – | – | – | R |
| FT012<sup>d</sup> | B1K-04-13 | ISR | 31.834361 | 35.304309 | JJ | 1 | 1 | – | – | – |
| FT013 | B1K-05-11 | ISR | 31.926131 | 35.469146 | JJ | 2 | 1 | – | 1 | R |
| FT017 | B1K-06-07 | ISR | 32.068642 | 35.398412 | JJ | 2 | – | 2 | – | R |
| FT021 | B1K-07-02 | ISR | 32.342164 | 35.515748 | JJ | – | – | – | – | R |
| FT041 | B1K-11-05 | ISR | 30.889786 | 34.631073 | NM | 1 | 1 | – | – | R |
| FT045 | B1K-12-06 | ISR | 31.672915 | 35.436719 | JJ | 1 | 1 | – | – | R |
| FT048 | B1K-13-01 | ISR | 32.852183 | 35.680149 | GH | – | – | – | – | R |
| FT057 | B1K-15-07 | ISR | 32.6906 | 35.662224 | GH | 2 | 2 | – | – | R |
| FT086 | B1K-21-08 | ISR | 32.647805 | 34.967268 | CG | 2 | 1 | 1 | – | S |
| FT087 | B1K-21-11 | ISR | 32.647599 | 34.967754 | CG | 3 | 2 | 1 | – | S |
| FT088 | B1K-21-20 | ISR | 32.647674 | 34.966946 | CG | 2 | 1 | 1 | – | S |
| FT113 | B1K-27-02 | ISR | 32.875386 | 35.544077 | HG | 2 | – | 1 | 1 | R |
| FT115 | B1K-27-11 | ISR | 32.874543 | 35.54338 | HG | 1 | 1 | – | – | R |
| FT120 | B1K-28-10 | ISR | 32.823909 | 35.497509 | HG | 1 | – | 1 | – | R |
| FT125 | B1K-29-20 | ISR | 32.821027 | 35.507253 | HG | 1 | – | – | 1 | R |
| FT132 | B1K-31-01 | ISR | 32.773355 | 35.44657 | NM | 2 | 2 | – | – | R |
| FT138 | B1K-32-08 | ISR | 32.578564 | 35.471275 | NM | 2 | 1 | 1 | – | S |
| FT145 | B1K-33-19 | ISR | 30.571722 | 34.676408 | HG | – | – | – | – | R |
| FT146 | B1K-34-04 | ISR | 30.845141 | 34.744956 | NM | 2 | 1 | – | 1 | R |
| FT152 | B1K-35-04 | ISR | 31.596864 | 34.898841 | SCJ | 1 | 1 | – | – | R |
| FT153<sup>e</sup> | B1K-35-08 | ISR | 31.596249 | 34.898955 | SCJ | 1<sup>e</sup> | 1 | –<sup>e</sup> | – | S |
| FT158 | B1K-36-05 | ISR | 32.925892 | 35.531067 | HG | 1 | – | – | 1 | R |
| FT169 | B1K-39-02 | ISR | 33.070336 | 35.769545 | GH | 1 | 1 | – | – | R |
| FT170 | B1K-39-20 | ISR | 33.070844 | 35.769422 | GH | 2 | 1 | 1 | – | S |
| FT174 | B1K-41-03 | ISR | 33.251503 | 35.647858 | SCJ | 2 | 2 | – | – | R |
| FT218 | B1K-49-18 | ISR | 31.771952 | 35.112916 | SCJ | 1 | – | – | 1 | R |
| FT231 | HID-4 | IRQ | 35.58333333 | 43 | UM | 1 | – | 1 | – | R |
| FT233 | HID-8-1 | IRQ | 36.41666667 | 41.65 | UM | 1 | – | 1 | – | R |
| FT248 | HID-53 | IRN | 31.67589444 | 48.57915556 | LM | 1 | – | – | 1 | R |
| FT256 | HID-65 | TUR | 36.75194444 | 37.47527778 | UM | 1 | – | 1 | – | R |
| FT271 | HID-104 | SYR | 36.77194444 | 40.86194444 | UM | 1 | – | 1 | – | R |
| FT281 | HID-137 | TUR | 36.85083333 | 40.0475 | UM | 2 | 1 | 1 | – | R |
| FT282 | HID-138 | IRN | 32 | 48.55 | LM | – | – | – | – | R |
| FT283 | HID-140 | IRQ | 36.4 | 44.2 | UM | 2 | 1 | 1 | – | R |
| FT287 | HID-145 | ISR | 31 | 34.91666667 | NM | 2 | 1 | 1 | – | S |
| FT289 | HID-149 | ISR | 32.96666667 | 35.6 | GH | 1 | – | 1 | – | R |
| FT293 | HID-160 | ISR | 31.6 | 34.9 | GH | 2 | 2 | – | – | R |
| FT313 | HID-213 | ISR | 31.4 | 34.6 | SCJ | 2 | 1 | – | 1 | R |
| FT320 | HID-230 | ISR | 32.25 | 34.85 | SCJ | 2 | 1 | 1 | – | R |
| FT349 | HID-302 | IRN | 32.55 | 48.81666667 | LM | 3 | 2 | 1 | – | R |
| FT352 | HID-307 | IRN | 31.53333333 | 48.81666667 | LM | 2 | 1 | 1 | – | R |
| FT353 | HID-308 | IRN | 31.53333333 | 48.81666667 | LM | 1 | 1 | – | – | R |
| FT355 | HID-310 | IRN | 31.53333333 | 48.81666667 | LM | 2 | – | 1 | 1 | R |
| FT393<sup>b</sup> | n.d. | n.d. | n.d. | n.d. | n.d. | 1 | 1 | – | – | R |
| FT394<sup>b</sup> | HID-386 | ISR | 31.7747333 | 35.0485083 | SCJ (mixed) | 1 | – | 1 | – | R |
| FT454 | HP-10-4 | TUR | 36.2558333 | 36.4672222 | NL | 1 | 1 | – | – | R |
| FT458 | HP-15-1 | TUR | 37.49333333 | 38.87666667 | NL | 1 | 1 | – | – | R |
| FT462 | HP-20-1 | TUR | 37.77833333 | 39.78 | NL | 2 | 1 | 1 | – | S |
| FT471 | HP-34-3 | TUR | 37.70222222 | 38.01222222 | NL | 1 | 1 | – | – | R |

<sup>a</sup> ISR = Israel, IRQ = Iraq, IRN = Iran, TUR = Turkey, and SYR = Syria; n.d. = not determined.
<sup>b</sup> FT393 and FT394 are also known as ISR42-8 and ISR101-23, respectively. Taxonomy of FT393 is not determined (n.d.).
<sup>c</sup> The reaction to the isolate ND90Pr of *B. sorokiniana*. R indicates resistant with disease rating 1-3 and S indicates susceptible with disease rating 5-9.
<sup>d</sup> FT012 was not tested for susceptibility to ND90Pr.
<sup>e</sup> Targeted resequencing identified a *Scs6* allele that escaped previous analysis, termed *FT153-2*.

**Table S6.**
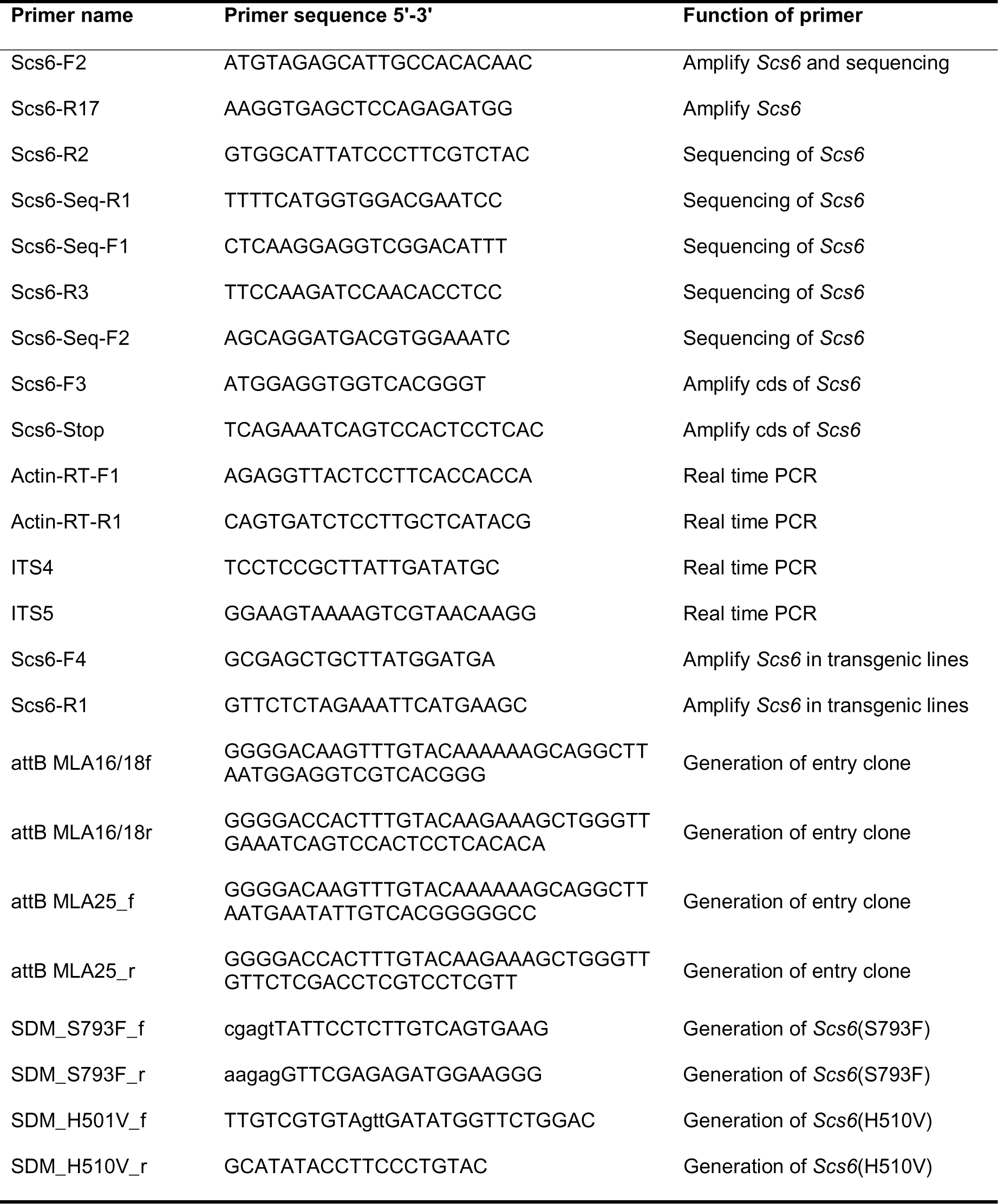
Primers used in this study.

